# The Repeating, Modular Architecture of the HtrA Proteases

**DOI:** 10.1101/2022.04.28.489847

**Authors:** Matthew Merski, Sandra Macedo-Ribeiro, Rafal M. Wieczorek, Maria W. Górna

## Abstract

A conserved, 26 residue sequence [AA(X_2_)[A/G][G/L](X_2_)GDV[I/L](X_2_)[V/L]NGE(X_1_)V(X_6_)] and corresponding structure repeating module was identified within the HtrA protease family using a non-redundant set (N=20) of publically available structures. While the repeats themselves were far from sequence perfect they had notable conservation to a statistically significant level with three or more repetitions identified within one protein at a level that would be expected to randomly occur only once per 1031 residues. This sequence repeat was associated with a six stranded antiparallel β-barrel module, two of which are present in the core of the structures of the PA clan of serine proteases, while a modified version of this module could be identified in the PDZ-like domains. Automated structural alignment methods had difficulties in superimposing these β-barrels but use of a target human HtrA2 structure showed that these modules had an average RMSD across the set of structures of less than 2 Å (mean and median). Our findings support Dayhoff’s hypothesis that complex proteins arose through duplication of simpler peptide motifs and domains.

## Introduction

Generally, the arrangement of amino acids in proteins is seemingly random (complex), although exceptions exist where notable patterns can be discerned in the amino acid sequence, such as low-complexity proteins^1^ or protein repeats^2-4^. Proteins also usually adopt distinct three dimensional structures and a wide variety of these have been reported in the public repository of the Protein Data Bank (PDB)^5^. A combination of elements (sequence, secondary structure, fold, and three-dimensional structure) comprise the architecture of the protein. However, the three-dimensional structure itself tends to be the most conserved aspect of the protein as both the sequence and function of the protein evolve much more quickly^6^, although it has been suggested that the folds themselves are fossils^7^ of the early, Archean proteins which may have evolved before the appearance of the last universal common ancestor (LUCA)^8^. Five decades ago based on the earliest protein structures, it was hypothesized that these primitive peptides formed oligomeric groups in solution which eventually fused into single peptide transcripts to give rise to the early, modern proteins which then over time gradually diverged into more complex forms^9-13^. This process of oligomerization followed by fusion has also been suggested to have given rise to repeat proteins, which are composed of a set of repeating structures and sequences 20-60 amino acids in length, which may have, over time evolved into complex, globular proteins (Fig. 1)^2,14,15^. While there are a number of well-known repeat protein types, they have generally received less researcher attention than globular proteins that have more complex structural architectures^16^ despite estimates suggesting that about a quarter of all known proteins have at least some repeat protein character^17^. This raises the obvious question as to what is obscuring the presence of all these expected protein repeats in structural databases such as the PDB, especially given the possibility that the early, ancestral proteins were all at least repeat-like^18-21^.

**Figure 1:**
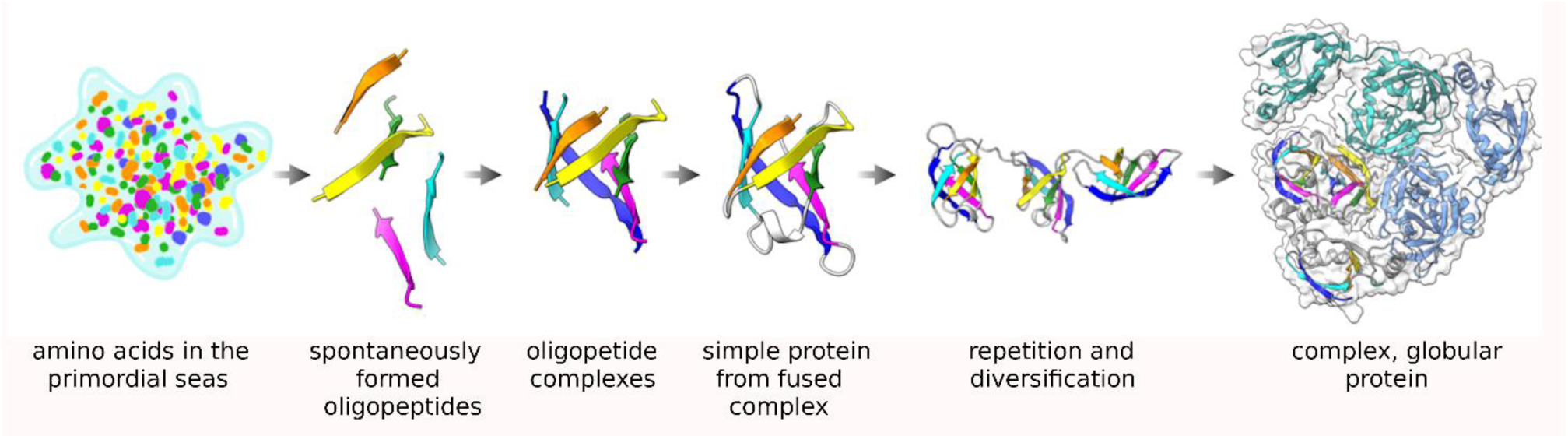
An illustration of Dayhoff’s hypothesis about the origin of proteins. From left to right, starting from individual, spontaneously formed amino acids in the Archaean seas, short oligopeptides formed spontaneously which then organized into homogenous complexes and eventually fused into a single transcript module, probably after being encoded in the genome. Duplication and repetition of these modules along with drift in their sequence and function eventually gave rise to complex, globular proteins.

Protein families that are widely distributed across the three kingdoms of life are likely to have roots deep in evolutionary time^8,14^, possibly even as far back as the Archean, pre-LUCA period and may be, in essence, representatives of such preserved, fossil architectures. One such protein family could be the HtrA family of proteases. The High Temperature Requirement A (HtrA) proteases are stress response, housekeeping proteases widely distributed throughout Nature^22^.

Notable examples of this family include DegP^23^ and DegS^24^ in prokaryotes, Deg1 in plants^25^, and HtrA2 in humans^26-29^. Structurally, HtrA proteases are members of the PA clan of serine proteases (including such notable examples as chymotrypsin A and thrombin) which contain a pair of six-stranded β-barrels^30^. HtrA proteases additionally have one or more C-terminal PDZ-like domains^27,31^. The PDZ domain is an 80-100 amino acid long protein interaction domain found in many different protein families^32^. In prokaryotes, DegP forms large 12- and 24-mer complexes while DegS exists as a simple trimer. In humans, the chromosomally encoded HtrA2 protease, linked to Parkinson’s disease^26^, functions as a housekeeping protease within the mitochondria^28^. Damage to the mitochondrial membranes results in leakage of HtrA2 into the cytoplasm where it digests peptide inhibitors of apoptosis leading to cell death^33^. HtrA2 has been shown to have an unusually high melting temperature^28^ and to preferably cleave unfolded substrate ensembles^34^. HtrA2 is maintained in a resting closed state and activated is a set of sequential steps that are initiated by binding of a hydrophobic motif to the PDZ-like domain, followed by exposure of the substrate binding site on the protease domain and activation of the proteolytic activity.

We have recently reported^35^ a survey of all the known protein sequences using a self-homology detection method based on DOTTER^36^. This allowed us to identify a number of protein families which had a notable amount of self-similarity, including the HtrA protease family. More detailed examination confirmed the initial detection of the repeating amino acid sequence and we were able to correlate the sequence repeats with a six strand antiparallel β-barrel structure that occurred at least three times in the monomeric structure of the protease, twice in the protease domain (a feature of the PA clan of serine proteases^30^) [Fig. 2] and once in each PDZ-like domain. These results suggest that the PDZ-like domain evolved from repetition of this basic barrel structure in the PA clan serine proteases.

**Figure 2:**
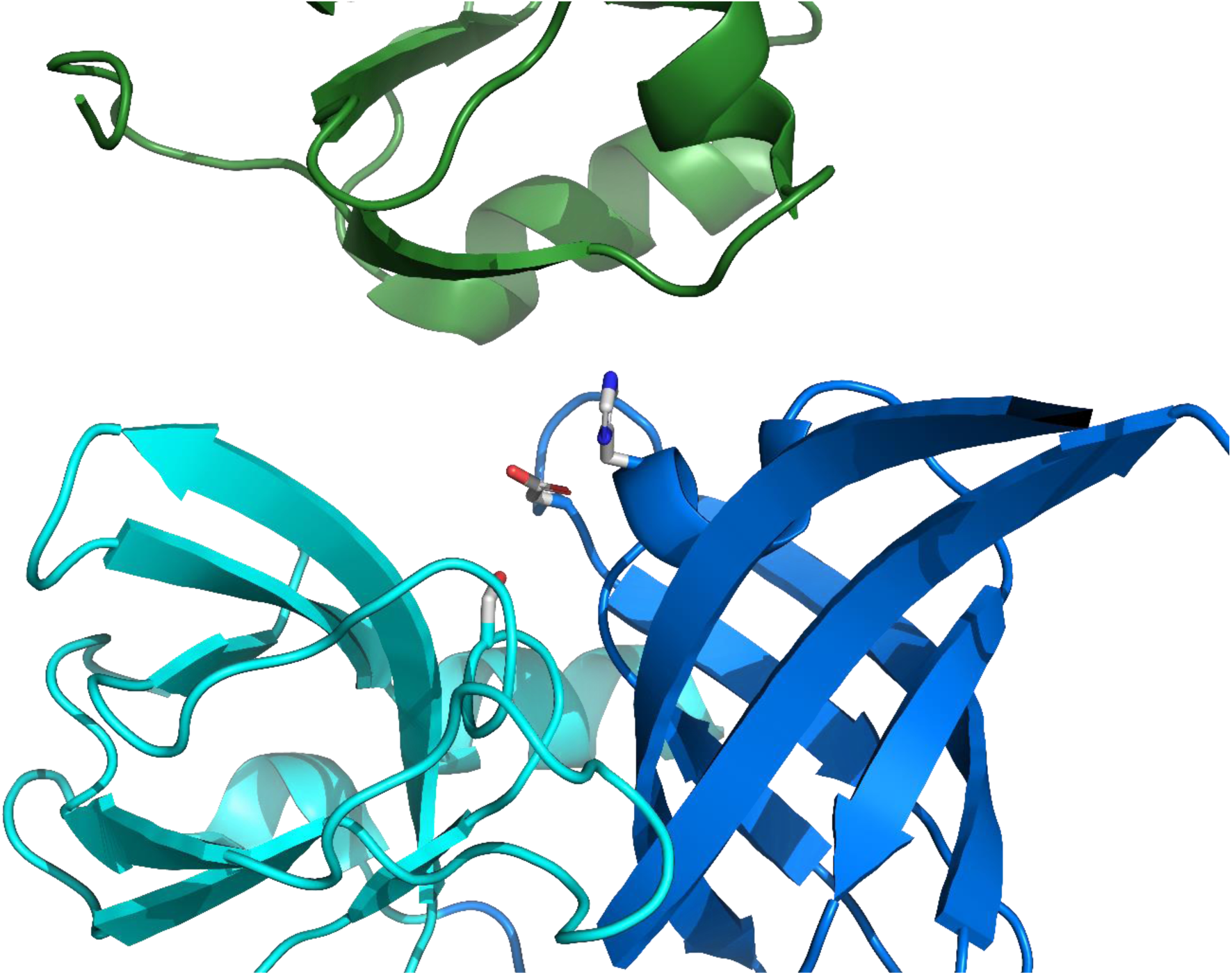
The active site in the HtrA proteases is separate between the modules. Cartoon diagram of human HtrA2 (PDB ID 5m3n^28^) showing the N-terminal protease (blue), C-terminal protease (cyan) and PDZ-like (green) modules. The catalytic triad of His198, Asp 228, and Ser306 are shown as sticks with white carbon, blue nitrogen, and red oxygen atoms.

## Results

Using a modified form of DOTTER to analyse protein self-homology^35^, the presence of notable self-homology was detected in the HtrA proteases. Reverse calculation of the homology plots produced by DOTTER^36^ using HtrA protease sequences from the PDB^5^ allowed the detection of a putative 26 amino acid repeat [AA(X_2_)[A/G][G/L](X_2_)GDV[I/L](X_2_)[V/L]NGE(X_1_)V(X_6_)] and alignment of the repeating regions with MUSCLE^37^ produced an initial estimate of the repeating sequence with 13 of the 26 positions being definable [Fig. 3, SI Fig. 1]. Comparison to random sequence and theoretical comparison to a Bernoulli model for cumulative probability [SI Figs. 2 & 3] suggested that high scoring matches (typically 5 or more matching positions) should be relatively uncommon (see SI methods) occurring randomly once every 1031 residues, which is much less than once per protein as the average HtrA protease monomer is 350-450 residues long^38^. Examination of the sequence unique HtrA proteases (90% ID) clearly identified three or more of these repeats in each monomer, corresponding roughly to two in the protease domain and one in each PDZ-like domain [SI Fig. 4] (some HtrA proteases such as DegP have two copies of the PDZ-like domain^32^). The protease and PDZ-like domains within the HtrA proteases do not have significant self-homology (mean = 18.2%, median = 15.3% for protease and PDZ-like 1 domains, N = 19 proteins [SI Table 1]). Only one protein, Legionella pneumophila DegQ (PDB ID 4ynn)^39^ had greater than 30% identity between its protease and PDZ-like domains (34.0 % ID). Low shared sequence identity has been previously noted in PDZ domains^40^. Further visual inspection of the structures identified a common alternating antiparallel six strand β-barrel module in the structures [Fig. 4, SI Fig. 5]. There is a shift in register in the PDZ-like module in which the first beta strand occurs outside of the barrel structure and the barrel remains unclosed as it contains only four additional strands [SI Fig 5]. The sixth strand is also rotated out of the structure or deleted, depending on the species. The alternating anti-parallel pattern present in the protease modules is maintained in the PDZ-like module but formally reversed as the designation of positive and negative strands is arbitrary. RMSD comparison of these isolated structures using PyMol found a poor structural similarity among the β-barrels with a mean RMSD of 4.1 Å between the two protease domains and mean RMSD values of 8.8 Å and 9.2 Å between the N-terminal protease module or the C-terminal protease module and the first PDZ-like domain, respectively [SI Table 2]. The superposition of the PDZ-like and protease modules was poor but their small size makes the RMSD values appear better than they are. Analysis by TM-align^41^ suggested good agreement between the protease domains (TM scores = 0.552 (mean), 0.549 (median) for the protease modules, 0.652 (mean), 0.638 (median) for the PDZ-like modules; RMSD = 2.80 Å (mean), 2.78 Å (median) for the protease domains, 2.28 Å (mean), 2.22 Å (median) for the PDZ-like domains). But, there was poor correspondence when the protease modules were compared to the PDZ-like modules or vice-versa (TM-score = 0.298 (mean), 0.299 (median); RMSD = 3.72 Å (mean), 3.70 Å (median)) [SI Table 3]. However, the modules from one human HtrA2 structure (PDB ID = 5m3n^28^) could be structurally aligned after manual examination [Fig. 4, SI Table 3]. The aligned modules from this specific PDB structure could then be used as “targets” for the corresponding modules in the other proteins. When this was done, the structural differences were generally minimized and good structural alignments could be achieved (mean = 2.9 Å, 2.9 Å, 1.9 Å; median = 2.5 Å, 2.8 Å, 1.7 Å for the N-terminal protease, C-terminal protease, and PDZ-like domain modules respectively) (SI Table 3).

**Figure 3:**
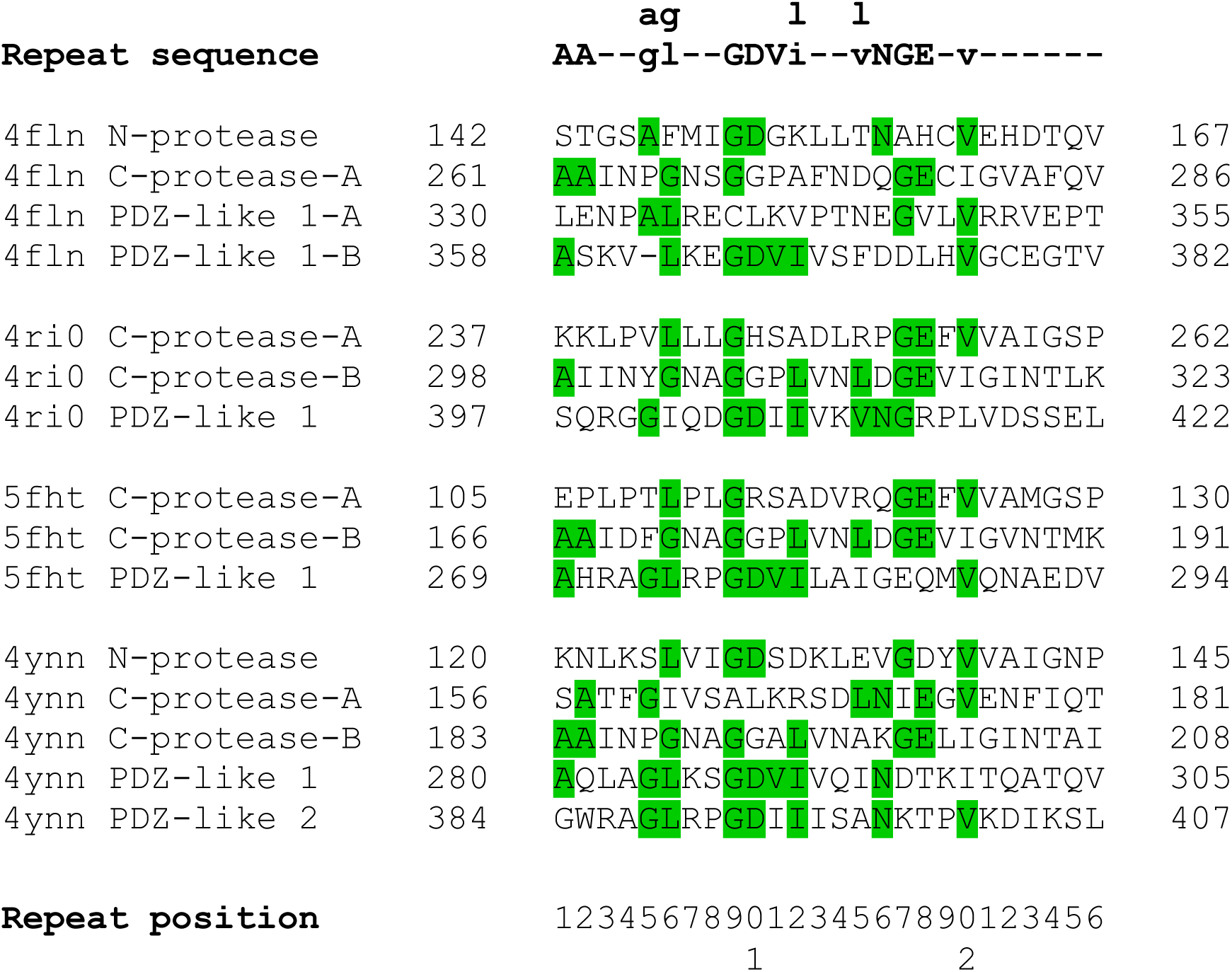
Identification of the sequence repeats in the HtrA proteases. The canonical sequence [AA(X_2_)[A/G][G/L](X_2_)GDV[I/L](X_2_)[V/L]NGE(X_1_)V(X_6_)] is shown on top in bold, with an additional residue shown for the four positions which have two possible canonical residues. Residues that match the canonical sequence are highlighted in green. The PDB id is given on the left along with the module the sequence is associated with. When a module has two copies of the sequence repeat, the first is denoted as A, the second as B. The beginning sequence position (using the PDB numbering) is to the left of the sequence while the ending position is given to the right of the sequence.

**Figure 4:**
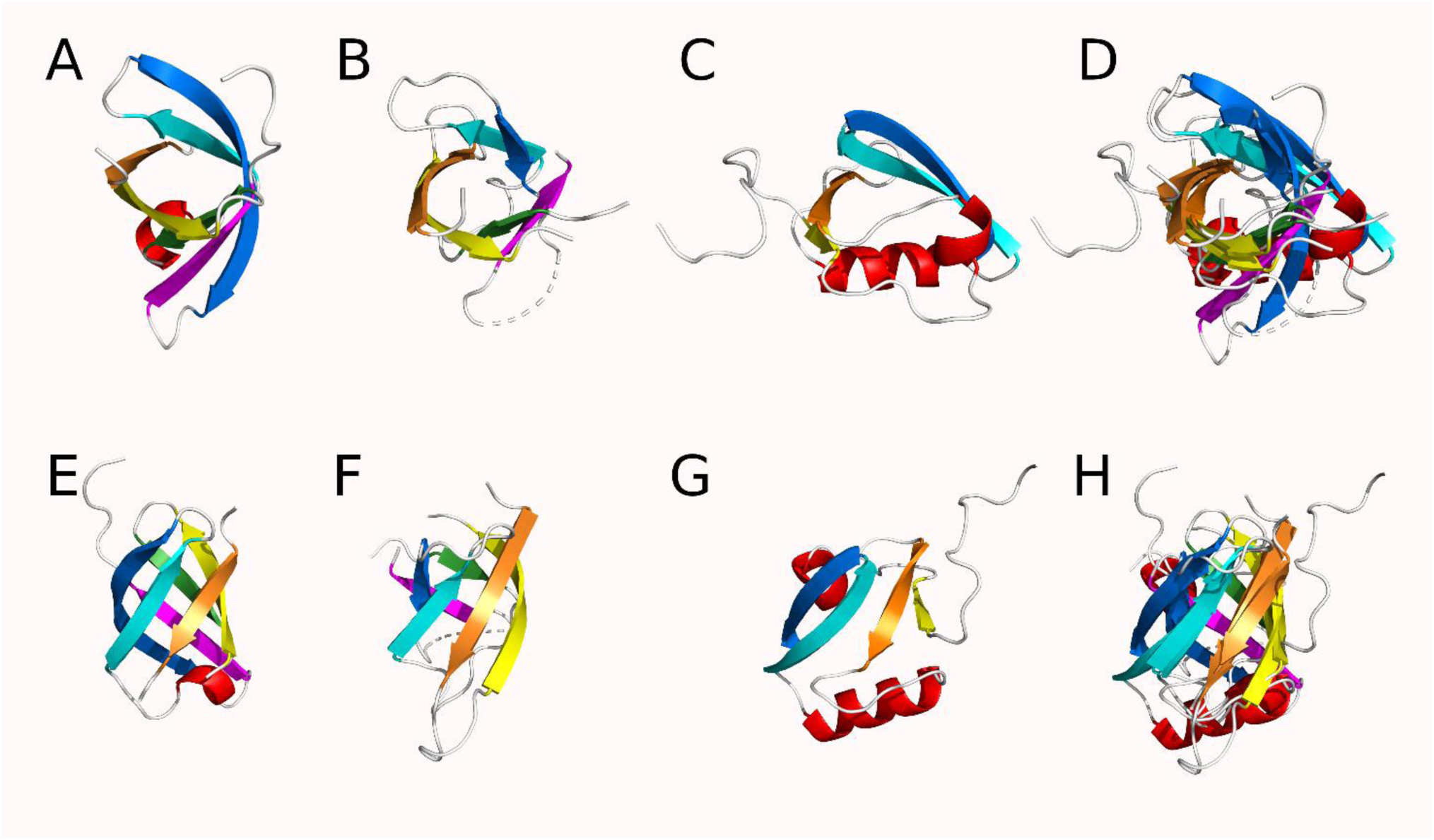
Cartoon diagram of the modules from HhoA, an HtrA protease from Synechocystis sp. PCC 6803 (PDB ID 5t69) showing the conserved structures of the HtrA modules. Strands are coloured orange, yellow, green, blue and magenta in order from N to C in the protease modules and in the equivalent spatial position in the PDZ-like module. Helices are coloured red and coil regions are white. A top-down view of the A) N-terminal protease module B) C-terminal protease module C) PDZ-like module and D) all three modules superimposed. Side views of the E) N-terminal protease module F) C-terminal protease module G) PDZ-like module and H) all three modules superimposed.

## Discussion

Fundamentally, the HtrA proteases are repeat proteins. A 26 residue sequence repeat associated with an anti-parallel β-barrel structure is clearly identifiable in the HtrA proteases [Figs. 2 & 4]. While the repeating sequences are far from perfect^42^, the frequency of matches to the defined canonical sequence is statistically significant [SI Figs. 2 & 3]. The individual modules can be structurally aligned to a set of target modules to a good average RMSD (< 3 Å) between the β-barrel structural modules within a given protein [Fig. 4, SI Table 3]. Repetitions of both sequence and structure in combination with the presence of two copies of a β-barrel in the PA clan of serine proteases^30^ (to which HtrA proteases belong) and that all three modules are have a peptide binding function strongly suggests that the HtrA proteases are the result of a set of repetitions of the ancestral β-barrel module followed by mutation and functional change in the third (by sequence order) module present in the PDZ-like domain(s)^43^.

The evolution of the modern HtrA protease structure from an ancestral β-barrel precursor, possibly an Archean, pre-LUCA protease^8^, via the PA clan ancestor^30^ offers an elegant solution to the problem of the origin of structural complexity of this family of proteases from a simple ancestor as suggested by Dayhoff’s hypothesis^9,18^. It is currently unknown if ancestral module itself was an active protease or if it simply had a peptide binding function and developed into an active protease after the duplication at the origin of the PA clan as the catalytic triad of the HtrA proteases is spread across the two protease modules. For example, in human HtrA2^27^, *E. coli* DegP^44^, and *A. thaliana* Deg2^45^, the catalytic serine is found in the C-terminal protease module, while the other two members of the triad are present in the first module. The PDZ-like module is likely derived from one of these modules as it contains several divergent structural features compared to the protease modules. The protease modules have six strands comprising its β-barrel, while only four of these are present in the PDZ-like domain module^43^ along with an additional N-terminal strand which is rotated out of the barrel structure [SI Fig. 5]. There are also many PA clan proteases which lack the PDZ-like domain which suggests that it evolved later. Therefore, while it is not undisputable, it seems likely that the PDZ-like module is a product of the duplication of one of the protease modules rather than the protease being derived from the PDZ-like domain. This may be an incorrect assumption, however, given the amount of lateral gene transfer that occurs in prokaryotes^46,47^.

By analysing the conserved self-homology patterns in the HtrA proteases, we were able to identify the simple, repeating β-barrel architecture present in this family. To the best of our knowledge, this repeating architecture has gone unremarked upon despite the fact that structures of these proteins have been publicly available for 20 years now^27,31^ and the widely recognized pair of β-barrels present in the PA clan of proteases. This was likely at least partially due to the general difficulty in identifying protein repeats^48-50^. In this specific case, there are one (or two) sequence repeats present in each of the β-barrel structural modules [SI Fig. 1], a small but notable discrepancy between the sequence and structural repetitions, which would contribute to the difficulty in identifying these repeats. However, a discrepancy between sequence and structural repeat is not uncommon in repeat proteins^51^. Additionally, automated structural alignment methods had difficult in detecting the similarity between the modules, even after the repeats had been unambiguously identified. The low sequence similarity between the members of the family or the different repeat modules as well as the low sequence conservation within the repeats themselves likely contributed to this detection issue, as did the variability in the assignment of the secondary structures in the protein structure models themselves [SI Tables 2 & 3]. It is also worth noting that standard structural alignments using PyMol failed to properly superimpose the protease modules with PDZ-like modules, even when they had been formally recognized although they could be convinced to superimpose the modules when a properly superimposed “bait” structure was used [SI table 3, SI Fig. 6].

Nevertheless, the successful detection of this overlooked repeat architecture in a well-studied family of proteins using self-homology does not imply that this method for repeat detection cannot be further improved. While the method was able to find the sequence repeats fairly easily, they did not correspond to the structural repeat and identification of that required significant human intervention. Even when the β-barrel was recognized, the movement of the first and last strand out of the barrel and the spatial rearrangement of the strands prevented accurate matching of the β-barrels using structural alignment algorithms without human optimization. Clearly, improvements in the automation of the structure search strategies would be beneficial here, since simple removal of the coil regions did not improve detection ability, quite likely due to discrepancies in identification of secondary structural features in the crystallographic models [SI Table 3]. And finally, despite identification of these repeating modules, it still cannot be indisputably determined which module is the most ancestral and which are derived.

Despite these caveats, the conserved, repeating architecture of the HtrA proteases is clearly identifiable in the family. Self-homology analysis was able to identify this architecture which had gone overlooked for decades, a clear success for this method of repeat detection. This repeat architecture shows an elegant method to generate complex protein structures from simple oligopeptide building blocks and might serve to inform protein engineering efforts. This repeat detection methodology can (and will) be applied to other well-studied protein families and potentially identify their underlying repeat architectures.

## Methods

A full description of the methods used here are included within the Supplemental Material.

## Acknowledgements

This work was supported by National Science Centre, Poland grant #2014/15/D/NZ1/00968 to M.W.G and by the European Union’s Horizon 2020 research and innovation programme under GA 952334 (PhasAGE) to S.M.-R. The authors would like to Dominik Gront for helpful discussions.

## References

1 Rado-Trilla, N. & Alba, M. M. Dissecting the role of low-complexity regions in the evolution of vertebrate proteins. Bmc Evol Biol 12 (2012).

2 Andrade, M. A., Perez-Iratxeta, C. & Ponting, C. P. Protein repeats: Structures, functions, and evolution. J Struct Biol 134, 117–131 (2001).

3 Espada, R. et al. Repeat proteins challenge the concept of structural domains. Biochem Soc T 43, 844–849 (2015).

4 Kajava, A. V. Tandem repeats in proteins: From sequence to structure. J Struct Biol 179, 279–288 (2012).

5 Burley, S. K. et al. RCSB Protein Data Bank: biological macromolecular structures enabling research and education in fundamental biology, biomedicine, biotechnology and energy. Nucleic Acids Res 47, D464–D474 (2019).

6 Illergard, K., Ardell, D. H. & Elofison, A. Structure is three to ten times more conserved than sequence-A study of structural response in protein cores. Proteins 77, 499–508 (2009).

7 Alva, V., Soding, J. & Lupas, A. N. A vocabulary of ancient peptides at the origin of folded proteins. Elife 4 (2015).

8 Ranea, J. A. G., Sillero, A., Thornton, J. M. & Orengo, C. A. Protein superfamily evolution and the last universal common ancestor (LUCA). J Mol Evol 63, 513–525 (2006).

9 Eck, R. V. & Dayhoff, M. O. Evolution of Structure of Ferredoxin Based on Living Relics of Primitive Amino Acid Sequences. Science 152, 363-& (1966).

10 Alva, V. & Lupas, A. N. From ancestral peptides to designed proteins. Curr Opin Struc Biol 48, 103–109 (2018).

11 Broom, A. et al. Modular Evolution and the Origins of Symmetry: Reconstruction of a Three-Fold Symmetric Globular Protein. Structure 20, 161–171 (2012).

12 Jackson, V. A., Busby, J. N., Janssen, B. J. C., Lott, J. S. & Seiradake, E. Teneurin Structures Are Composed of Ancient Bacterial Protein Domains. Front Neurosci-Switz 13 (2019).

13 Wieczorek, R. On Prebiotic Ecology, Supramolecular Selection and Autopoiesis. Origins Life Evol B 42, 445–450 (2012).

14 Lupas, A. N., Ponting, C. P. & Russell, R. B. On the evolution of protein folds: Are similar motifs in different protein folds the result of convergence, insertion, or relics of an ancient peptide world? J Struct Biol 134, 191–203 (2001).

15 Soding, J. & Lupas, A. N. More than the sum of their parts: on the evolution of proteins from peptides. Bioessays 25, 837–846 (2003).

16 Paladin, L. et al. RepeatsDB 2.0: improved annotation, classification, search and visualization of repeat protein structures (vol 45, pg D308, 2016). Nucleic Acids Res 45, 3613–3613 (2017).

17 Marcotte, E. M., Pellegrini, M., Yeates, T. O. & Eisenberg, D. A census of protein repeats. J Mol Biol 293, 151–160 (1999).

18 Romero, M. L. R., Rabin, A. & Tawfik, D. S. Functional Proteins from Short Peptides: Dayhoff’s Hypothesis Turns 50. Angew Chem Int Edit 55, 15966–15971 (2016).

19 Laurino, P. et al. An Ancient Fingerprint Indicates the Common Ancestry of Rossmann-Fold Enzymes Utilizing Different Ribose-Based Cofactors. Plos Biol 14 (2016).

20 Galpern, E. A., Freiberger, M. I. & Ferreiro, D. U. Large Ankyrin repeat proteins are formed with similar and energetically favorable units. Plos One 15 (2020).

21 Zhu, H. B. et al. Origin of a folded repeat protein from an intrinsically disordered ancestor. Elife 5 (2016).

22 Clausen, T., Kaiser, M., Huber, R. & Ehrmann, M. HTRA proteases: regulated proteolysis in protein quality control. Nat Rev Mol Cell Bio 12, 152–162 (2011).

23 Ortega, J., Iwanczyk, J. & Jomaa, A. Escherichia coli DegP: a Structure-Driven Functional Model. J Bacteriol 191, 4705–4713 (2009).

24 Sohn, J., Grant, R. A. & Sauer, R. T. Allostery Is an Intrinsic Property of the Protease Domain of DegS IMPLICATIONS FOR ENZYME FUNCTION AND EVOLUTION. J Biol Chem 285, 34039–34047 (2010).

25 Kley, J. et al. Structural adaptation of the plant protease Deg1 to repair photosystem II during light exposure. Nat Struct Mol Biol 18, 728–731 (2011).

26 Hegde, R. et al. Identification of Omi/HtrA-2 as a mitochondrial apoptotic serine protease that disrupts inhibitor of apoptosis protein-caspase interaction. J Biol Chem 277, 432–438 (2002).

27 Li, W. Y. et al. Structural insights into the pro-apoptotic function of mitochondrial serine protease HtrA2/Omi. Nat Struct Biol 9, 436–441 (2002).

28 Merski, M. et al. Molecular motion regulates the activity of the Mitochondrial Serine Protease HtrA2. Cell Death Dis 8 (2017).

29 Zurawa-Janicka, D. et al. Temperature-induced changes of HtrA2(Omi) protease activity and structure. Cell Stress Chaperon 18, 35–51 (2013).

30 Di Cera, E. Serine Proteases. Iubmb Life 61, 510–515 (2009).

31 Krojer, T., Garrido-Franco, M., Huber, R., Ehrmann, M. & Clausen, T. Crystal structure of DegP (HtrA) reveals a new protease-chaperone machine. Nature 416, 455–459 (2002).

32 Lee, H. J. & Zheng, J. J. PDZ domains and their binding partners: structure, specificity, and modification. Cell Commun Signal 8 (2010).

33 Martins, L. M. et al. Binding specificity and regulation of the serine protease and PDZ domains of HtrA2/Omi. J Biol Chem 278, 49417–49427 (2003).

34 Toyama, Y., Harkness, R. W. & Kay, L. E. Structural basis of protein substrate processing by human mitochondrial high-temperature requirement A2 protease. P Natl Acad Sci USA 119, e2203172119. doi:doi.org/10.1073/pnas.2203172119 (2022).

35 Merski, M. et al. Self-analysis of repeat proteins reveals evolutionarily conserved patterns. Bmc Bioinformatics 21 (2020).

36 Sonnhammer, E. L. L. & Durbin, R. A dot-matrix program with dynamic threshold control suited for genomic DNA and protein sequence analysis (Reprinted from Gene Combis, vol 167, pg GC1-GC10, 1996). Gene 167, Gc1–Gc10 (1995).

37 Edgar, R. C. MUSCLE: multiple sequence alignment with high accuracy and high throughput. Nucleic Acids Res 32, 1792–1797 (2004).

38 Clausen, T., Southan, C. & Ehrmann, M. The HtrA family of proteases: Implications for protein composition and cell fate. Mol Cell 10, 443–455 (2002).

39 Schubert, A., Wrase, R., Hilgenfeld, R. & Hansen, G. Structures of DegQ from Legionella pneumophila Define Distinct ON and OFF States. J Mol Biol 427, 2840–2851 (2015).

40 Teyra, J., Ernst, A., Singer, A., Sicheri, F. & Sidhu, S. S. Comprehensive analysis of all evolutionary paths between two divergent PDZ domain specificities. Protein Sci 29, 433–442 (2020).

41 Zhang, Y. & Skolnick, J. TM-align: a protein structure alignment algorithm based on the TM-score. Nucleic Acids Res 33, 2302–2309 (2005).

42 Jorda, J., Xue, B., Uversky, V. N. & Kajava, A. V. Protein tandem repeats - the more perfect, the less structured. Febs J 277, 2673–2682 (2010).

43 Skelton, N. J. et al. Origins of PDZ domain ligand specificity - Structure determination and mutagenesis of the erbin PDZ domain. J Biol Chem 278, 7645–7654 (2003).

44 Krojer, T. et al. Structural basis for the regulated protease and chaperone function of DegP. Nature 453, 885–U831 (2008).

45 Sun, R. H. et al. Crystal Structure of Arabidopsis Deg2 Protein Reveals an Internal PDZ Ligand Locking the Hexameric Resting State. J Biol Chem 287, 37564–37569 (2012).

46 Burroughs, A. M., Allen, K. N., Dunaway-Mariano, D. & Aravind, L. Evolutionary genomics of the HAD superfamily: Understanding the structural adaptations and catalytic diversity in a superfamily of phosphoesterases and allied enzymes. J Mol Biol 361, 1003–1034 (2006).

47 Vos, M., Hesselman, M. C., te Beek, T. A., van Passel, M. W. J. & Eyre-Walker, A. Rates of Lateral Gene Transfer in Prokaryotes: High but Why? Trends Microbiol 23, 598–605 (2015).

48 Schaper, E., Kajava, A. V., Hauser, A. & Anisimova, M. Repeat or not repeat?-Statistical validation of tandem repeat prediction in genomic sequences. Nucleic Acids Res 40, 10005–10017 (2012).

49 Marold, J. D., Kavran, J. M., Bowman, G. D. & Barrick, D. A Naturally Occurring Repeat Protein with High Internal Sequence Identity Defines a New Class of TPR-like Proteins. Structure 23, 2055–2065 (2015).

50 Gul, I. S., Hulpiau, P., Saeys, Y. & van Roy, F. Metazoan evolution of the armadillo repeat superfamily. Cell Mol Life Sci 74, 525–541 (2017).

51 Renault, L. et al. The 1.7 angstrom crystal structure of the regulator of chromosome condensation (RCC1) reveals a seven-bladed propeller. Nature 392, 97–101 (1998).

