## Supplemental Material for "The Repeating, Modular Architecture of the HtrA Proteases"

**Methods:** As previously reported, all known proteins (UniRef90[1]) were examined for self-homology using a modified version of DOTTER[2,3]. This analysis found a number of proteins with significant self-homology that were in a protease Do-like cluster. HtrA proteases were then collected from the PDB[4] and short sequences (less than 200 residues) were removed and the remainder were filtered at the 90% sequence identity threshold with CD-HIT[5] leaving 20 unique structures (*i.e.* 2zle, 2z9i, 3gdv, 3nzi, 3pv5, 3qo6, 4a9g, 4fln, 4ic5, 4ic6, 4ri0, 4ynn, 5fht, 5ilb, 5jyk, 5t69, 5zvj, 6jjo, 6z05, 7co3). The locations of probable sequence repeats were identified by reverse calculation of the DOTTER plots of these protein sequences where each residue was assigned its maximal self-homology score. The high scoring regions from all the proteins were separated and the frequency of each amino acid at each position was calculated and those positions which had strong biases towards a single, or a pair of similar, amino acids were noted. This process identified a 26 residue repeating sequence. Multiple sequence alignment of these repeats with MUSCLE[6] helped to clarify the repeated sequence in which 13 of the 26 positions were conserved, specifically [AA($X_2$)[A/G][G/L]($X_2$)GDV[I/L]($X_2$)[V/L]NGE($X_1$)V($X_6$)] which can also be represented as AA--[A/G][G/L]--GDV[I/L]--[V/L]NGE-V------. This repeat was then used for further sequence searches.

The statistical significance of this sequence repeat was verified by comparison of the repeat pattern to randomly generated sequence. A score estimate for the random sequence was defined by a binomial (Bernoulli) model. The random chance for success at each position was defined as the probability for the expected amino acid at that positon for each of the 13 defined positions. More explicitly, suppose a set of coin flips (Bernoulli trials) such that each event will have 13 trials and each of those trials have a probability of success defined by the natural frequency of the amino acids that are acceptable in that position. The score for each event is defined by the number of successes that occur in the event. For each success a score of 1 / P(x) where P(x) is the probability of finding an acceptable amino acid in that position is given while failures get a score of 1. For positions with 2 acceptable amino acids, the probability is the sum of the two amino acid frequencies. This allowed a success score to be defined for any actual sequence as:

$$Score = \Pi(T)$$

where the score value for success T(S) in any position is:

$$T(S) = 1/P\ (x)$$

and for failure T(F):

$$T(F) = 1$$

Failures are given a score of one in order to not modify the score (*i.e.* a sequence with no matches to the defined repeat sequence received a score of 1). To further verify this model, a 9996060 residue length of random amino acid sequence was generated by the Sequence Manipulation Suite[7] and each 26 residue sequence unit was compared to the repeat sequence to generate an estimate of the probability for the repeat sequence to appear randomly (SI Figure 2 & 3). Sequences with a score of greater than 90,000 appeared with a frequency of less than 1 per 1000 residues, in agreement with the theoretical binomial model.

The repeat sequence was then identified in a multiple sequence alignment (MUSCLE) in the set of protein structures (SI Fig. 1). The sequence repeats did not share a common secondary structure. However, a shared common antiparallel β-barrel structure could be identified within the structures, which was associated with the sequence repeats (SI Fig. 5). These were comprised of a pair of six stranded barrels in the protease domain and a partial, four stranded barrel in each PDZ-like domain (SI Fig. 4). The PDB structures were divided into the individual barrel structures and compared by structural alignment in PyMol[8]. While generally, the alignment of the domains to each other was poor using PyMol, the three domains from a specific human HtrA2 protease (PDB ID 5m3n) could all be superimposed when aligned in PyMol. The N terminal protease, C terminal protease, and PDZ domain modules from all the other example HtrA proteases could then be aligned to the equivalent module from the 5m3n structure and a good superimposition was achieved (SI Fig.6). The modules were also compared by TM-align[9].

**Supplemental References:**

1 Bateman, A. *et al.* UniProt: a worldwide hub of protein knowledge. *Nucleic Acids Res* **47**, D506-D515 (2019).

2 Sonnhammer, E. L. L. & Durbin, R. A dot-matrix program with dynamic threshold control suited for genomic DNA and protein sequence analysis (Reprinted from Gene Combis, vol 167, pg GC1-GC10, 1996). *Gene* **167**, Gc1-Gc10 (1995).

3 Merski, M. *et al.* Self-analysis of repeat proteins reveals evolutionarily conserved patterns. *Bmc Bioinformatics* **21** (2020).

4 Burley, S. K. *et al.* RCSB Protein Data Bank: biological macromolecular structures enabling research and education in fundamental biology, biomedicine, biotechnology and energy. *Nucleic Acids Res* **47**, D464-D474 (2019).

5 Huang, Y., Niu, B. F., Gao, Y., Fu, L. M. & Li, W. Z. CD-HIT Suite: a web server for clustering and comparing biological sequences. *Bioinformatics* **26**, 680-682 (2010).

6 Madeira, F. *et al.* The EMBL-EBI search and sequence analysis tools APIs in 2019. *Nucleic Acids Res* **47**, W636-W641 (2019).

7 Stothard, P. The sequence manipulation suite: JavaScript programs for analyzing and formatting protein and DNA sequences. *Biotechniques* **28**, 1102-+ (2000).

8 The PyMOL Molecular Graphics System.

9 Zhang, Y. & Skolnick, J. TM-align: a protein structure alignment algorithm based on the TM-score. *Nucleic Acids Res* **33**, 2302-2309 (2005).

**<u>SI Figure 1:</u> MUSCLE 3.8 alignment of HtrA proteases**

Multiple sequence alignment of unique HtrA sequences from the PDB. The β-barrel structures from the PDB structures are identified with light grey highlighting (SI Figure 4). Potential sequence repeat regions are identified by dark grey highlighting while residues matching the canonical repeat sequence are shown with green highlighting. However, structure 4a9g lacked explicit identification of its secondary structures within the PDB model and PDB ID 3nzi lacked a modeled PDZ-like domain so those β-barrel modules were not indicated with grey highlighting.

```
4fln            --------------------------NAESSNPPQKMAFKAFG-----------------
5ilb            --------------------------GHDASF----------------------------
5jyk            -------------MGSSHH-----HHHHSSGLVPRGSHMASMTGGQQMGRGSEFELLNNE
5fht            ------------------------------MAVP--------------------------
3nzi            -------------MG---------------------------------------------
4ri0            ------------------------------------------------------------
6z05            ------------------------------------------------------------
2z9i            -----------------------------ANMPP--------------------------
5zvj            ------------------------MVDAFTTSKVT-------------------------
3gdv            -------------MRGSHH-----HHHHGRSLNPLS------------------------
7co3            MPKALRFLGWPVLVGVLLALLIIQHNPELVGLPRQEVHVEQ-------------------
3pv5            -------------MRGSHH-----HHHHGSAEPPN-------------------------
4ynn            -------------MRGSHH-----HHHHGSAQDLT-------------------------
4a9g            -----------------------SIPGQVADQAP--------------------------
2zle            ------------------------------------------------------------
6jjo            -------------MGSSHH-----HHHHSSGLVPRGSHMA--------------------
5t69            ---------------------GIDPFTMADDLPPAPVI----------------------
4ic5            -------------MGSSHH-----HHHHSSGLVPRGSHMASAL-----------------
4ic6            -------------MGLGDP-----SVATVEDVSPT-------------------------
3qo6            -------------------------MAAFVVSTPK-------------------------

4fln            -----------SPKKEKKESLSDFSRDQQTDPAKIHDASFLNAVVKVYC-----------
5ilb            ----------------------------------------LNAVVKVYC-----------
5jyk            SEAGNQRTSSPERSRSRLHHSDTKNGDCSNGMIVSTTTESIPAAPSWETVVKVVPSMDAV
5fht            --------------SPPPASPRSQYNFIADVVEKT-----APAVVYIEI-----------
3nzi            --------------QEDPNSLRHKYNFIADVVEKI-----APAVVHIEL-----------
4ri0            --------------LHQLSSPRYKFNFIADVVEKI-----APAVVHIEL-----------
6z05            --------------VLSYHDS----------IKDA-----KKSVVNISTSKTITRANRPS
2z9i            -------------------------GSVEQVAAKV-----VPSVVMLET-----------
5zvj            --------------LSTTGNAQEPAGRFTKVAAAV-----ADSVVTIES-----------
3gdv            --------------TPQFDSTDETPASYNLAVRRA-----APAVVNVYN-----------
7co3            --------------APLLSRLQEGPVSYANAVSRA-----APAVANLYTTKMVSKPSHPL
3pv5            --------------MPSMAPV----------LKNI-----MPAIVNVAVQGYLP------
4ynn            -----------------------NMPSLAPVLKNA-----MPAIVNVAVQGYLPNNM---
4a9g            --------------LPSLAPM----------LEKV-----LPAVVSVRVEGTASQGQK--
2zle            -------------AETSSATTAQQMPSLAPMLEKV-----MPSVVSINVEGSTTVNTPR-
6jjo            --------------ETSSATTAQQMPSLAPMLEKV-----MPSVVSINVEGSTTVNTPR-
5t69            --------------TAQASVPLTSESFVAAAVSRS-----GPAVVRIDTETVVTRRTDPI
4ic5            -----------EQFKEKEEELEEEEERNVNLFQKT-----SPSVVYIEA-----------
4ic6            --------------VFPAGPLFPTEGRIVQLFEKN-----TYSVVNIFD-----------
3qo6            --------------KLQTDEL-----ATVRLFQEN-----TPSVVYITN-----------
                                                          :
```

```
4fln   ----------THTAPDYSLPWQKQ----------------------RQFTSTGSAFMIG-
5ilb   ----------THTAPDYSLPWQKQ----------------------RQFTSTGSAFMIG-
5jyk   V-----KVFCVHTEPNFSLPWQRK----------------------RQYSSGSSGFIIG-
5fht   ---------LDRHPFLGREVPI----------------------------SNGSGFVVAA
3nzi   ---------FRKLPFSKREVPV----------------------------ASGSGFIVSE
4ri0   ---------FLRHPLFGRNVPL----------------------------SSGSGFIMSE
6z05   PLDDFFNDPYFKQFFDFDFPQRKGKNDK------------------EVVSSLGSGVIISK
2z9i   -------------DLGRQSEE------------------------------GSGIILSA
5zvj   --------------VSDQEGM------------------------------QGSGVIVDG
3gdv   ---------RGLNTNSHNQLEIR---------------------------TLGSGVIMDQ
7co3   F-----DDPMFRRFFGDNLPQQK-----------------------RMESSLGSAVIMSA
3pv5   ------NDVTPPGSAGNDEENQPNNRPPQSRMPEKG----------RKFESIGSGVIIDP
4ynn   ------ASGNADDDDGENSKQPSRIPEKG-----------------RKFESIGSGVIIDP
4a9g   ------IPEEFKKFFGDDLPDQPA----------------------QPFEGLGSGVIINA
2zle   ------MPRNFQQFFGDDSPFCQEGSPFQSSPFCQGGQGGNGGGQQQKFMALGSGVIIDA
6jjo   ------MPRNFQQFFGDDSPFCQEGSPFQSSPFCQGGQGGNGGGQQQKFMALGSGVIIDA
5t69   L-----DDPFFQEFFGRSFPVPPRE---------------------RRIAGQGSGFIIDN
4ic5   ------IEL-PKTSSGDILTDEEN----------------------GKIEGTGSGFVWDK
4ic6   ------VTLRPQLKMTGVVEIP---------------------------EGNGSGVVWDG
3qo6   ------LAVRQDAFTLDVLEVP---------------------------QGSGSGFVWDK
                                                            .*..:

4fln   -DGKLLTNAHCVEHD--T---------QVKVKRRGDDR-KYVAKVLVRGVDCDIALLSVE
5ilb   -DGKLLTNAHCVEHD--T---------QVKVKRRGDDR-KYVAKVLVRGVDCDIALLSVE
5jyk   -GRRVLTNAHSVEHH--T---------QVKLKKRGSDT-KYLATVLAIGTECDIALLTVT
5fht   -DGLIVTNAHVVADR--R---------RVRVRLLSG-D-TYEAVVTAVDPKADIATLRIQ
3nzi   -DGLIVTNAHVVTNK--H---------RVKVELKNG-A-TYEAKIKDVDEKADIALIKID
4ri0   -AGLIITNAHVVSSN--SAAPGRQ---QLKVQLQNG-D-SYEATIKDIDKKSDIATIKIH
6z05   -DGYIVTNNHVVDDA--D---------TITVNLPGSDI-EYKAKLIGKDPKTDLAVIKIE
2z9i   -EGLILTNNHVIAAA--AKPPLGSPPPKTTVTFSDG-R-TAPFTVVGADPTSDIAVVRVQ
5zvj   -RGYIVTNNHVISEA--ANNPSQF---KTTVVFNDG-K-EVPANLVGRDPKTDLAVLKVD
3gdv   -RGYIITNKHVINDA--D---------QIIVALQDG-R-VFEALLVGSDSLTDLAVLKIN
7co3   -EGYLLTNNHVTAGA--D---------QIIVALRDG-R-ETIAQLVGSDPETDLAVLKID
3pv5   NNGVIITNDHVIRNA--S---------LITVTLQDG-R-RLKARLIGGDSETDLAVLKID
4ynn   KNGIIITNDHVIRNA--N---------LITVTLQDG-R-RLKARLIGGDSETDLAVLKID
4a9g   SKGYVLTNNHVINQA--Q---------KISIQLNDG-R-EFDAKLIGSDDQSDIALLQIQ
2zle   DKGYVVTNNHVVDNA--T---------VIKVQLSDG-R-KFDAKMVGKDPRSDIALIQIQ
6jjo   DKGYVVTNNHVVDNA--T---------VIKVQLSDG-R-KFDAKMVGKDPRSDIALIQIQ
5t69   -SGIILTNAHVVDGA--S---------KVVVTLRDG-R-TFDGQVRGTDEVTDLAVVKIE
4ic5   -LGHIVTNYHVIAKL--ATDQFGLQRCKVSLVDAKGTRFSKEGKIVGLDPDNDLAVLKIE
4ic6   -QGYIVTNYHVIGNALSRNPSPGDVVGRVNILASDGVQKNFEGKLVGADRAKDLAVLKVD
3qo6   -QGHIVTNYHVIRGA--S---------DLRVTLADQ-T-TFDAKVVGFDQDKDVAVLRID
           ::** *                    :             :   .   *:* : :
```

```
4fln    SEDFWKGAEPLRLGHLPRLQDSVTVVGYPLGG----DTISVTKGVVSRIEVT-----SYA
5ilb    SEDFWKGAEPLRLGHLPRLQDSVTVVGYPLGG----DTISVTKGVVSRIEVT-----SYA
5jyk    DDEFWEGVSPVEFGDLPALQDAVTVVGYPIGG----DTISVTSGVVSRMEIL-----SYV
5fht    TKEP-LPTLPLGRSADVRQGEFVVAMGSPFAL-----QNTITSGIVSSAQRPAR---DLG
3nzi    HQGK-LPVLLLGRSSELRPGEFVVAIGSPFSL-----QNTVTTGIVSTTQRGGK---ELG
4ri0    PKKK-LPVLLLGHSADLRPGEFVVAIGSPFAL-----QNTVTTGIVSTAQREGR---ELG
6z05    ANN--LSAITFTNSDDLMEGDVVFALGNPFGV-----GFSVTSGIISALNK------DNI
2z9i    GVSG-LTPISLGSSSDLRVGQPVLAIGSPLGL-----EGTVTTGIVSALNRPVSTTGEAG
5zvj    NVDN-LTVARLGDSSKVRVGDEVLAVGAPLGL-----RSTVTQGIVSALHRPVPLSGEGS
3gdv    ATGG-LPTIPINARRVPHIGDVVLAIGNPYNL-----GQTITQGIISATGR------IGL
7co3    LKN--LPAMTLGRSDGIRTGDVCLAIGNPFGV-----GQTVTMGIISATGR------NQL
3pv5    AKN--LKSLVIGDSDKLEVGDFVVAIGNPFGLNSFGNSQSATFGIVSALKR------SDL
4ynn    AKN--LKSLVIGDSDKLEVGDYVVAIGNPFGLNSFGNSQSATFGIVSALKR------SDL
4a9g    NPSK-LTQIAIADSDKLRVGDFAVAVGNPFGL-----GQTATSGIVSALGR------SGL
2zle    NPKN-LTAIKMADSDALRVGDYTVAIGNPFGL-----GETVTSGIVSALGR------SGL
6jjo    NPKN-LTAIKMADSDALRVGDYTVAIGNPFGL-----GETVTSGIVSALGR------SGL
5t69    PQGSALPVAPLGTSSNLQVGDWAIAVGNPVGL-----DNTVTLGIISTLGRSAA---QAG
4ic5    TEGRELNPVVLGTSNDLRVGQSCFAIGNPYGY-----ENTLTIGVVSGLGREI----PSP
4ic6    APETLLKPIKVGQSNSLKVGQQCLAIGNPFGF-----DHTLTVGVISGLNRDI----FSQ
3qo6    APKNKLRPIPVGVSADLLVGQKVFAIGNPFGL-----DHTLTTGVISGLRREIS---SAA
                  .         :   .:* *          : * *::*

4fln    HGSSDLLGIQIDAAINPGNSGGPAFNDQGECIGVAFQV--YRSEETE------NIGYVIP
5ilb    HGSSDLLGIQIDAAINPGNSGGPAFNDQGECIGVAFQV--YRSEETE------NIGYVIP
5jyk    HGSTELLGLQIDAAINSGNSGGPAFNDKGKCVGIAFQS--LKHEDAE------NIGYVIP
5fht    LPQTNVEYIQTDAAIDFGNAGGPLVNLDGEVIGVNTMK--VTA----------GISFAIP
3nzi    LRNSDMDYIQTDAIINYGNSGGPLVNLDGEVIGINTLK--VTA----------GISFAIP
4ri0    LRDSDMDYIQTDAIINYGNAGGPLVNLDGEVIGINTLK--VTA----------GISFAIP
6z05    GLNQYENFIQTDASINPGNSGGALVDSRGYLVGINSAI--LSR--GGG---NNGIGFAIP
2z9i    NQNTVLDAIQTDAAINPGNSGGALVNMNAQLVGVNSAIATLGADSADAQSGSIGLGFAIP
5zvj    DTDTVIDAIQTDASINHGNAGGPLIDMDAQVIGINTAG--KSLSDSAS-----GLGFAIP
3gdv    NPTGRQNFLQTDASINPGNSGGALVNSLGELMGINTLS--FDKSNDGE--TPEGIGFAIP
7co3    GLNTYEDFIQTDAAINPGNAGGALVDAAGNLIGINTAI--FSK--SGG---SQGIGFAIP
3pv5    NIEGVENFIQTDAAIGGGNSGGALVNAKGELIGINTAI--LSP--YGG---NVGIGFAIP
4ynn    NIEGVENFIQTDAAINPGNAGGALVNAKGELIGINTAI--ISP--YGG---NVGIGFAIP
4a9g    NLEGLENFIQTDASINRGNAGGALLNLNGELIGINTAI--LAP--GGG---SVGIGFAIP
2zle    NAENYENFIQTDAAINRGNSGGALVNLNGELIGINTAI--LAP--DGG---NIGIGFAIP
6jjo    NAENYENFIQTDAAINRGNAGGALVNLNGELIGINTAI--LAP--DGG---NIGIGFAIP
5t69    IPDKRVEFIQTDAAINPGNAGGPLLNARGEVIGINTAI--RAD--------ATGIGFAIP
4ic5    NGKSISEAIQTDADINSGNAGGPLLDSYGHTIGVNTAT--FTRKGSGM---SSGVNFAIP
4ic6    TGVTIGGGIQTDAAINPGNAGGPLLDSKGNLIGINTAI--FTQ--TGT---SAGVGFAIP
3qo6    TGRPIQDVIQTDAAINPGNSGGPLLDSSGTLIGINTAI--YSP--SGA---SSGVGFSIP
                :* ** *. **:**. .:  .  :*:                   .:.: **
```

```
4fln                TTVVSHFLTD------YERNGKYTGYPCLGVLLQKLENPALR-----ECLKVP---TN-E
5ilb                TTVVSHFLTD------YERNGKYTGFPVLGIEWQKMENPDLR-----KSMGME---SHQK
5jyk                TPVIVHFIQD------YEKHDKYTGFPVLGIEWQKMENPDLR-----KSMGME---SHQK
5fht                SDRLREFLHRGEKKNSSSGISGSQ-RRYIGVMMLTL-SPSILAELQLREPSFP---DVQH
3nzi                SDKIKKFLTESHDRQ-AKGKAITK-KKYIGIRMMSL-TSSKAKELKDRHRDFP---DVIS
4ri0                SDRITRFLTEFQ----DKQIKDWK-KRFIGIRMRTI-TPSLVDELKASNPDFP---EVSS
6z05                SNMVKDIAKK------LIEKGKID-RGFLGVTILAL-QGDTK-----KAYKNQ------E
2z9i                VDQAKRIADE------LISTGKAS-HASLGVQVT---NDKDT-----------------L
5zvj                VNEMKLVANS------LIKDGKIV-HPTLGISTRSV-SNAIA-----------------S
3gdv                FQLATKIMDK------LIRDGRVI-RGYIGIGGREI-APLHA-----QGGGID---QL-Q
7co3                TKLALEVMQS------IIEHGQVI-RGWLGVEVKAL-TPELA-----ESLGLG---ET-A
3pv5                INMVKDVAQQ------IIKFGSIH-RGLMGIFVQHL-TPELA-----QAMGYP---EDFQ
4ynn                INMAKDVAQQ------IIKFGSIH-RGLMGIFVQHL-TPELA-----QSMGYA---EDFQ
4a9g                SNMARTLAQQ------LIDFGEIK-RGLLGIKGTEM-SADIA-----KAFNLD---VQ-R
2zle                SNMVKNLTSQ------MVEYGQVK-RGELGIMGTEL-NSELA-----KAMKVD---AQ-R
6jjo                SNMVKNLTSQ------MVEYGQVK-RGELGIMGTEL-NSELA-----KAMKVD----AQR
5t69                IDQAKAIQNT------LAAGGTVP-HPYIGVQMMNI-TVDQA-QQNNRNPNSPFIIPEVD
4ic5                IDTVVRTVPY------LIVYGTAY-RDRLSS------VDKLA------------------
4ic6                SSTVLKIVPQ------LIQFSKVL-RAGINIELA---PDPVA-----NQLNVR------N
3qo6                VDTVGGIVDQ------LVRFGKVT-RPILGIKF----APDQS----VEQLGV-------S
                                                    :.

4fln                GVLVRRVEPTSDASKV-LKE-----------GDVIVSFDDLHVGCEGTVPFRSSERIAFR
5ilb                GVRIRRIEPTAPESQV-LKP-----------SDIILSFDGVNIANDGTVPFRHGERIGFS
5jyk                GVRIRRIEPTAPESQV-LKP-----------SDIILSFDGVNIANDGTVPFRHGERIGFS
5fht                GVLIHKVILGSPAHRAGLRP-----------GDVILAIGEQMV----------QNAEDVY
3nzi                GAYIIEVIPDTPAEAGGLKE-----------NDVIISINGQSV----------VSANDVS
4ri0                GIYVQEVAPNSPSQRGGIQD-----------GDIIVKVNGRPL----------VDSSELQ
6z05                GALITDVQKGSSADEAGLKR-----------GDLVTKVNNKVI----------KSPIDLK
2z9i                GAKIVEVVAGGAAANAGVPK-----------GVVVTKVDDRPI----------NSADALV
5zvj                GAQVANVKAGSPAQKGGILE-----------NDVIVKVGNRAV----------ADSDEFV
3gdv                GIVVNEVSPDGPAANAGIQV-----------NDLIISVDNKPA----------ISALETM
7co3                GIVVAGVYRDGPAARGGLLP-----------GDVILTIDKQEA----------SDGRRSM
3pv5                GALVSQVNPNSPAELAGLKA-----------GDIITQINDTKI----------TQATQVK
4ynn                GALVSQVNQNSPAQLAGLKS-----------GDVIVQINDTKI----------TQATQVK
4a9g                GAFVSEVLPGSGSAKAGVKA-----------GDIITSLNGKPL----------NSFAELR
2zle                GAFVSQVLPNSSAAKAGIKA-----------GDVITSLNGKPI----------SSFAALR
6jjo                GAFVSQVLPNSSAAKAGIKA-----------GDVITSLNGKPI----------SSFAALR
5t69                GILVMRVLPGTPAERAGIRR-----------GDVIVAVDGTPI----------SDGARLQ
4ic5                ------------------------------------------------------------
4ic6                GALVLQVPGKSLAEKAGLHPTSRGFAGNIVLGDIIVAVDDKPV----------KNKAELM
3qo6                GVLVLDAPPSGPAGKAGLQSTKRDGYGRLVLGDIITSVNGTKV----------SNGSDLY
```

```
4fln     YLISQKFAGDIAEIGII-RAGEHKKVQVVLRPRVHLVPYHIDGGQPSYIIVAGLVFTPLS
5ilb     YLISQKYTGDSALVKVL-RNKEILEFNIKLAIHKRLIPAHISGKPPSYFIVAGFVFTTVS
5jyk     YLISQKYTGDSALVKVL-RNKEILEFNIKLAIHKRLIPAHISGKPPSYFIVAGFVFTTVS
5fht     EAVRT---QSQLAVQIR-RGRETLTLYVTPEVTE------LEHHHHHH------------
3nzi     DVIKR---ESTLNMVVR-RGNEDIMITVIPEEID---PRSLEHHHHHH------------
4ri0     EAVLT---ESPLLLEVR-RGNDDLLFSIAPEVVMG------GHHHHHH------------
6z05     NYIGTLEIGQKISLSYE-RDGENKQASFILKGEKENPKGVQS--DLIDGLSLRNLDPRLK
2z9i     AAVRSKAPGATVALTFQDPSGGSRTVQVTLGKAEQ-----LEHHHHHH------------
5zvj     VAVRQLAIGQDAPIEVV-REGRHVTLTVKPDPDSTKLAAALEHHHHHH------------
3gdv     AQVAEIRPGSVIPVVVM-RDDKQLTLQVTIQEYPATN-----------------------
7co3     NQVARTRPGQKISIVVL-RNGQKVNLTAEVGLRPPPAPAPQQKQDGGE------------
3pv5     TTISLLRVGSTVKIIVE-RDNKPLTLSAVVTDIKSHEQKLQSNNPFLYGLALRAFEQESP
4ynn     TTISLLRAGSTAKIKIL-RDNKPLTLDVEVTDIKKHEQKLQSNNPFLYGLALRNFEQESP
4a9g     SRIATTEPGTKVKLGLL-RNGKPLEVEVTLDTSTSSSASAEMITPALEGATLSDGQL---
2zle     AQVGTMPVGSKLTLGLL-RDGKQVNVNLELQQSSQNQVDSSSIFNGIEGAEMSN------
6jjo     AQVGTMPVGSKLTLGLL-RDGKQVNVNLELQQSSQNQVDSSSIFNGIEGAEMSN------
5t69     RIVEQAGLNKALKLDLL-RGDRRLSLTVQTAQLRNPTS----------------------
4ic5     --------------------------------------AALEHHHHHH------------
4ic6     KILDEYSVGDKVTLKIK-RGNEDLELKISLEEKSS-----LEHHHHHH------------
3qo6     RILDQCKVGDEVTVEVL-RGDHKEKISVTLEPKPDESAAALEHHHHHH------------

4fln     EPLIE----EECEDTIGLKLLTKARYSVARFRGEQIVILSQVLANEVNIGYEDMNNQQVL
5ilb     VPYLRSEYGKEYEFDAPVKLLEKHLHAMAQSVDEQLVVVSQVLVSDINIGYEEIVNTQVV
5jyk     VPYLRSEYGKEYEFDAPVKLLEKHLHAMAQSVDEQLVVVSQVLVSDINIGYEEIVNTQVV
5fht     ------------------------------------------------------------
3nzi     ------------------------------------------------------------
4ri0     ------------------------------------------------------------
6z05     DRLQI-------------------------PKDVNGVLVDSVKEKSKGKNSGFQEGDIII
2z9i     ------------------------------------------------------------
5zvj     ------------------------------------------------------------
3gdv     ------------------------------------------------------------
7co3     ------------------------------------------------------------
3pv5     P-----------------------------HGNVIGVQVVGASENSAGWRAGIRPGDIII
4ynn     P-----------------------------HGNVVGVQVVGASETSAGWRAGLRPGDIII
4a9g     ------------------------------KDGGKGIKIDEVVKGSPAAQAGLQKDDVII
2zle     ------------------------------KGKDQGVVVNNVKTGTPAAQIGLKKGDVII
6jjo     ------------------------------KGKDQGVVVNNVKTGTPAAQIGLKKGDVII
5t69     ------------------------------------------------------------
4ic5     ------------------------------------------------------------
4ic6     ------------------------------------------------------------
3qo6     ------------------------------------------------------------
```

```
4fln          KFNGIPIRNIHHLAHLIDMCKDKYLVFEFEDNYVAVLEREASNSASLCILKDYGIPSERS
5ilb          AFNGKPVKNLKGLAGMVENCEDEYMKFNLDYDQIVVLDTKTAKEATLDILTTHCIPSAMS
5jyk          AFNGKPVKNLKGLAGMVENCEDEYMKFNLDYDQIVVLDTKTAKEATLDILTTHCIPSAMS
5fht          ------------------------------------------------------------
3nzi          ------------------------------------------------------------
4ri0          ------------------------------------------------------------
6z05          GVGQSEIKNLKDLEQALKQVNKKEFTKVWVYRNGFATLLVLK------------------
2z9i          ------------------------------------------------------------
5zvj          ------------------------------------------------------------
3gdv          ------------------------------------------------------------
7co3          ------------------------------------------------------------
3pv5          SANKKPVTDVKSLQTIAQEKKKELLVQVLRGPGSMYLLVI--------------------
4ynn          SANKTPVKDIKSLQAVAHEKGKQLLVQVLRGAGALYLLII--------------------
4a9g          GVNRDRVNSIAEMRKVLAAKPAIIALQIVRGNESIYLLMRLEHHHHHH------------
2zle          GANQQAVKNIAELRKVLDSKPSVLALNIQRGDSTIYLLMQ--------------------
6jjo          GANQQAVKNIAELRKVLDSKPSVLALNIQRGDSTIYLLMQ--------------------
5t69          ------------------------------------------------------------
4ic5          ------------------------------------------------------------
4ic6          ------------------------------------------------------------
3qo6          ------------------------------------------------------------

4fln          ADLLEPYVDPIDDTQALDQGIGDSPVSNLEIGFDGLVWA
5ilb          DDLKTEERN------------------------------
5jyk          DDLKTEERN------------------------------
5fht          ---------------------------------------
3nzi          ---------------------------------------
4ri0          ---------------------------------------
6z05          ---------------------------------------
2z9i          ---------------------------------------
5zvj          ---------------------------------------
3gdv          ---------------------------------------
7co3          ---------------------------------------
3pv5          ---------------------------------------
4ynn          ---------------------------------------
4a9g          ---------------------------------------
2zle          ---------------------------------------
6jjo          ---------------------------------------
5t69          ---------------------------------------
4ic5          ---------------------------------------
4ic6          ---------------------------------------
3qo6          ---------------------------------------
```

**<u>SI Figure 2:</u> Frequency of HtrA Repeats in Random Protein Sequences**

Comparison of the scoring of random protein sequences to the HtrA repeat sequence. All the randomly generated possible sequences were scored according to the Bernoulli statistical model and the scores were binned. The fraction of sequences that occurred in each bin is plotted against the minimum score for that bin. Note that both axes are log based. A vertical red line indicates the score cutoff used to identify HtrA repeat sequences within the sequence unique PDB set.

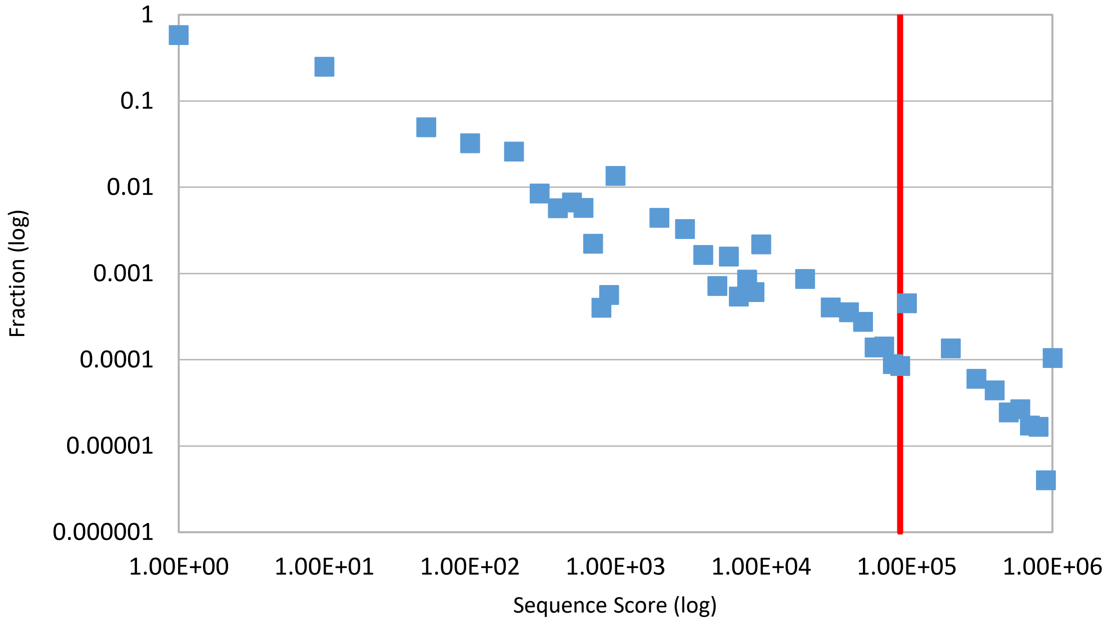

**<u>SI Figure 3:</u> Expected protein length of a random sequence matching that score to occur**

The expected length of a protein in which a sequence scoring X or higher should occur randomly, as extrapolated from randomly generated protein sequences. That is, a protein of length N would be expected to have a section of sequence with that scores at least as high as X somewhere within its length. Note the log scale of the y-axis. A vertical red line indicates the score cutoff used to identify HtrA repeat sequences, the same score as in SI Fig. 2. The intersection of this red line and the data points indicates the expected length of a protein sequence needed for a random occurrence of a sequence scoring at least that high to occur within that protein, in this case 1031 residues.

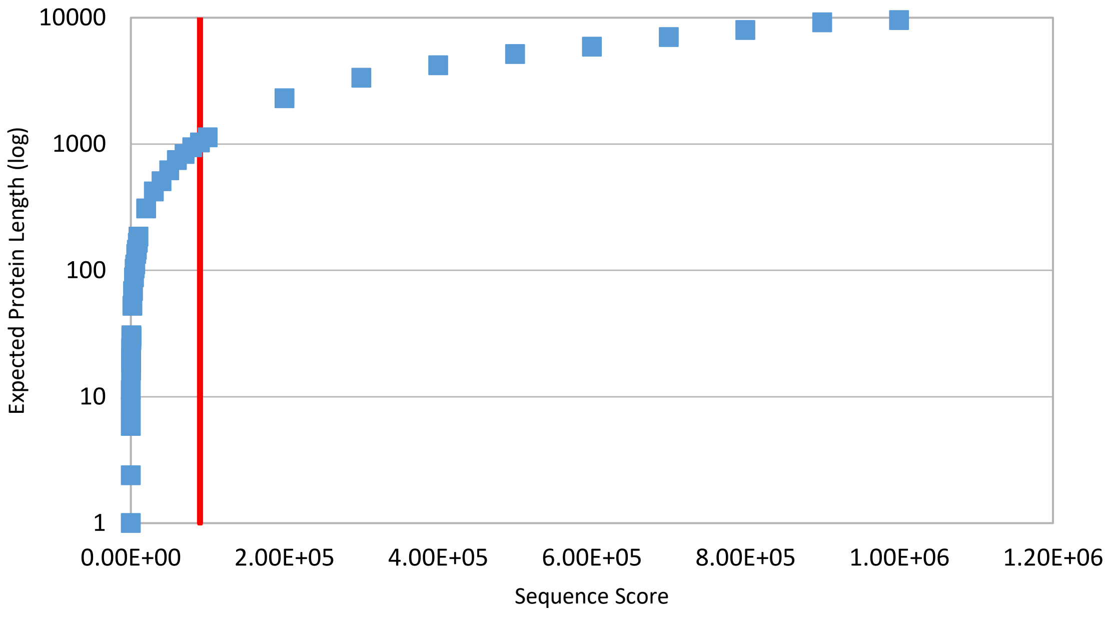

**<u>SI Figure 4:</u> Identification of the β-barrel structures in the HtrA proteases**

The conserved β-barrel modules are explicitly identified by highlighting. The N-terminal protease module in cyan, the C-terminal protease module in magenta, PDZ-like 1 and PDZ-like 2 in orange and violet respectively.

```
CLUSTAL multiple sequence alignment by MUSCLE (3.8)
4FLN            --------------------------NAESSNPPQKMAFKAFG-----------------
5ILB            --------------------------GHDASF----------------------------
5JYK            -------------MGSSHH-----HHHHSSGLVPRGSHMASMTGGQQMGRGSEFELLNNE
5FHT            ------------------------------MAVP--------------------------
3NZI            -------------MG---------------------------------------------
4RI0            ------------------------------------------------------------
6Z05            ------------------------------------------------------------
2Z9I            -----------------------------ANMPP--------------------------
5ZVJ            ------------------------MVDAFTTSKVT-------------------------
3GDV            -------------MRGSHH-----HHHHGRSLNPLS------------------------
7CO3            MPKALRFLGWPVLVGVLLALLIIQHNPELVGLPRQEVHVEQ-------------------
3PV5            -------------MRGSHH-----HHHHGSAEPPN-------------------------
4YNN            -------------MRGSHH-----HHHHGSAQDLT-------------------------
4A9G            -----------------------SIPGQVADQAP--------------------------
2ZLE            ------------------------------------------------------------
6JJO            -------------MGSSHH-----HHHHSSGLVPRGSHMA--------------------
5T69            ---------------------GIDPFTMADDLPPAPVI----------------------
4IC5            -------------MGSSHH-----HHHHSSGLVPRGSHMASAL-----------------
4IC6            -------------MGLGDP-----SVATVEDVSPT-------------------------
3QO6            -------------------------MAAFVVSTPK-------------------------

4FLN            -----------SPKKEKKESLSDFSRDQQTDPAKIHDASFLNAVVKVYC-----------
5ILB            ----------------------------------------LNAVVKVYC-----------
5JYK            SEAGNQRTSSPERSRSRLHHSDTKNGDCSNGMIVSTTTESIPAAPSWETVVKVVPSMDAV
5FHT            --------------SPPPASPRSQYNFIADVVEKT-----APAVVYIEI-----------
3NZI            --------------QEDPNSLRHKYNFIADVVEKI-----APAVVHIEL-----------
4RI0            --------------LHQLSSPRYKFNFIADVVEKI-----APAVVHIEL-----------
6Z05            --------------VLSYHDS----------IKDA-----KKSVVNISTSKTITRANRPS
2Z9I            -------------------------GSVEQVAAKV-----VPSVVMLET-----------
5ZVJ            --------------LSTTGNAQEPAGRFTKVAAAV-----ADSVVTIES-----------
3GDV            --------------TPQFDSTDETPASYNLAVRRA-----APAVVNVYN-----------
7CO3            --------------APLLSRLQEGPVSYANAVSRA-----APAVANLYTTKMVSKPSHPL
3PV5            --------------MPSMAPV----------LKNI-----MPAIVNVAVQGYLP------
4YNN            -----------------------NMPSLAPVLKNA-----MPAIVNVAVQGYLPNNM---
4A9G            --------------LPSLAPM----------LEKV-----LPAVVSVRVEGTASQGQK--
2ZLE            -------------AETSSATTAQQMPSLAPMLEKV-----MPSVVSINVEGSTTVNTPR-
6JJO            --------------ETSSATTAQQMPSLAPMLEKV-----MPSVVSINVEGSTTVNTPR-
5T69            --------------TAQASVPLTSESFVAAAVSRS-----GPAVVRIDTETVVTRRTDPI
4IC5            -----------EQFKEKEEELEEEEERNVNLFQKT-----SPSVVYIEA-----------
4IC6            --------------VFPAGPLFPTEGRIVQLFEKN-----TYSVVNIFD-----------
3QO6            --------------KLQTDEL-----ATVRLFQEN-----TPSVVYITN-----------
                                                          :
```

```
4FLN            ----------THTAPDYSLPWQKQ----------------------RQFTSTGSAFMIG-
5ILB            ----------THTAPDYSLPWQKQ----------------------RQFTSTGSAFMIG-
5JYK            V-----KVFCVHTEPNFSLPWQRK----------------------RQYSSGSSGFIIG-
5FHT            --------LDRHPFLGREVPI-----------------------------SNGSGFVVAA
3NZI            --------FRKLPFSKREVPV-----------------------------ASGSGFIVSE
4RI0            --------FLRHPLFGRNVPL-----------------------------SSGSGFIMSE
6Z05            PLDDFFNDPYFKQFFDFDFPQRKGKNDK------------------EVVSSLGSGVIISK
2Z9I            -------------DLGRQSEE-------------------------------GSGIILSA
5ZVJ            --------------VSDQEGM------------------------------QGSGVIVDG
3GDV            --------RGLNTNSHNQLEIR----------------------------TLGSGVIMDQ
7CO3            F-----DDPMFRRFFGDNLPQQK-----------------------RMESSLGSAVIMSA
3PV5            ------NDVTPPGSAGNDEENQPNNRPPQSRMPEKG----------RKFESIGSGVIIDP
4YNN            ------ASGNADDDDGENSKQPSRIPEKG-----------------RKFESIGSGVIIDP
4A9G            ------IPEEFKKFFGDDLPDQPA----------------------QPFEGLGSGVIINA
2ZLE            ------MPRNFQQFFGDDSPFCQEGSPFQSSPFCQGGQGGNGGGQQQKFMALGSGVIIDA
6JJO            ------MPRNFQQFFGDDSPFCQEGSPFQSSPFCQGGQGGNGGGQQQKFMALGSGVIIDA
5T69            L-----DDPFFQEFFGRSFPVPPRE---------------------RRIAGQGSGFIIDN
4IC5            ------IEL-PKTSSGDILTDEEN----------------------GKIEGTGSGFVWDK
4IC6            ------VTLRPQLKMTGVVEIP---------------------------EGNGSGVVWDG
3QO6            ------LAVRQDAFTLDVLEVP---------------------------QGSGSGFVWDK
                                                                    .*..:

4FLN            -DGKLLTNAHCVEHD--T---------QVKVKRRGDDR-KYVAKVLVRGVDCDIALLSVE
5ILB            -DGKLLTNAHCVEHD--T---------QVKVKRRGDDR-KYVAKVLVRGVDCDIALLSVE
5JYK            -GRRVLTNAHSVEHH--T---------QVKLKKRGSDT-KYLATVLAIGTECDIALLTVT
5FHT            -DGLIVTNAHVVADR--R---------RVRVRLLSG-D-TYEAVVTAVDPKADIATLRIQ
3NZI            -DGLIVTNAHVVTNK--H---------RVKVELKNG-A-TYEAKIKDVDEKADIALIKID
4RI0            -AGLIITNAHVVSSN--SAAPGRQ---QLKVQLQNG-D-SYEATIKDIDKKSDIATIKIH
6Z05            -DGYIVTNNHVVDDA--D---------TITVNLPGSDI-EYKAKLIGKDPKTDLAVIKIE
2Z9I            -EGLILTNNHVIAAA--AKPPLGSPPPKTTVTFSDG-R-TAPFTVVGADPTSDIAVVRVQ
5ZVJ            -RGYIVTNNHVISEA--ANNPSQF---KTTVVFNDG-K-EVPANLVGRDPKTDLAVLKVD
3GDV            -RGYIITNKHVINDA--D---------QIIVALQDG-R-VFEALLVGSDSLTDLAVLKIN
7CO3            -EGYLLTNNHVTAGA--D---------QIIVALRDG-R-ETIAQLVGSDPETDLAVLKID
3PV5            NNGVIITNDHVIRNA--S---------LITVTLQDG-R-RLKARLIGGDSETDLAVLKID
4YNN            KNGIIITNDHVIRNA--N---------LITVTLQDG-R-RLKARLIGGDSETDLAVLKID
4A9G            SKGYVLTNNHVINQA--Q---------KISIQLNDG-R-EFDAKLIGSDDQSDIALLQIQ
2ZLE            DKGYVVTNNHVVDNA--T---------VIKVQLSDG-R-KFDAKMVGKDPRSDIALIQIQ
6JJO            DKGYVVTNNHVVDNA--T---------VIKVQLSDG-R-KFDAKMVGKDPRSDIALIQIQ
5T69            -SGIILTNAHVVDGA--S---------KVVVTLRDG-R-TFDGQVRGTDEVTDLAVVKIE
4IC5            -LGHIVTNYHVIAKL--ATDQFGLQRCKVSLVDAKGTRFSKEGKIVGLDPDNDLAVLKIE
4IC6            -QGYIVTNYHVIGNALSRNPSPGDVVGRVNILASDGVQKNFEGKLVGADRAKDLAVLKVD
3QO6            -QGHIVTNYHVIRGA--S---------DLRVTLADQ-T-TFDAKVVGFDQDKDVAVLRID
                    ::** *                    :             :   .   *:* : :

4FLN            SEDFWKGAEPLRLGHLPRLQDSVTVVGYPLGG----DTISVTKGVVSRIEVT-----SYA
5ILB            SEDFWKGAEPLRLGHLPRLQDSVTVVGYPLGG----DTISVTKGVVSRIEVT-----SYA
5JYK            DDEFWEGVSPVEFGDLPALQDAVTVVGYPIGG----DTISVTSGVVSRMEIL-----SYV
5FHT            TKEP-LPTLPLGRSADVRQGEFVVAMGSPFAL-----QNTITSGIVSSAQRPAR---DLG
3NZI            HQGK-LPVLLLGRSSELRPGEFVVAIGSPFSL-----QNTVTTGIVSTTQRGGK---ELG
4RI0            PKKK-LPVLLLGHSADLRPGEFVVAIGSPFAL-----QNTVTTGIVSTAQREGR---ELG
6Z05            ANN--LSAITFTNSDDLMEGDVVFALGNPFGV-----GFSVTSGIISALNK------DNI
2Z9I            GVSG-LTPISLGSSSDLRVGQPVLAIGSPLGL-----EGTVTTGIVSALNRPVSTTGEAG
5ZVJ            NVDN-LTVARLGDSSKVRVGDEVLAVGAPLGL-----RSTVTQGIVSALHRPVPLSGEGS
3GDV            ATGG-LPTIPINARRVPHIGDVVLAIGNPYNL-----GQTITQGIISATGR------IGL
7CO3            LKN--LPAMTLGRSDGIRTGDVCLAIGNPFGV-----GQTVTMGIISATGR------NQL
3PV5            AKN--LKSLVIGDSDKLEVGDFVVAIGNPFGLNSFGNSQSATFGIVSALKR------SDL
4YNN            AKN--LKSLVIGDSDKLEVGDYVVAIGNPFGLNSFGNSQSATFGIVSALKR------SDL
4A9G            NPSK-LTQIAIADSDKLRVGDFAVAVGNPFGL-----GQTATSGIVSALGR------SGL
2ZLE            NPKN-LTAIKMADSDALRVGDYTVAIGNPFGL-----GETVTSGIVSALGR------SGL
6JJO            NPKN-LTAIKMADSDALRVGDYTVAIGNPFGL-----GETVTSGIVSALGR------SGL
5T69            PQGSALPVAPLGTSSNLQVGDWAIAVGNPVGL-----DNTVTLGIISTLGRSAA---QAG
4IC5            TEGRELNPVVLGTSNDLRVGQSCFAIGNPYGY-----ENTLTIGVVSGLGREI----PSP
4IC6            APETLLKPIKVGQSNSLKVGQQCLAIGNPFGF-----DHTLTVGVISGLNRDI----FSQ
3QO6            APKNKLRPIPVGVSADLLVGQKVFAIGNPFGL-----DHTLTTGVISGLRREIS---SAA
                          .         :   .:* *          : * *::*
```

```
4FLN        HGSSDLLGIQIDAAINPGNSGGPAFNDQGECIGVAFQV--YRSEETE------NIGYVIP
5ILB        HGSSDLLGIQIDAAINPGNSGGPAFNDQGECIGVAFQV--YRSEETE------NIGYVIP
5JYK        HGSTELLGLQIDAAINSGNSGGPAFNDKGKCVGIAFQS--LKHEDAE------NIGYVIP
5FHT        LPQTNVEYIQTDAAIDFGNAGGPLVNLDGEVIGVNTMK--VTA----------GISFAIP
3NZI        LRNSDMDYIQTDAIINYGNSGGPLVNLDGEVIGINTLK--VTA----------GISFAIP
4RI0        LRDSDMDYIQTDAIINYGNAGGPLVNLDGEVIGINTLK--VTA----------GISFAIP
6Z05        GLNQYENFIQTDASINPGNSGGALVDSRGYLVGINSAI--LSR--GGG---NNGIGFAIP
2Z9I        NQNTVLDAIQTDAAINPGNSGGALVNMNAQLVGVNSAIATLGADSADAQSGSIGLGFAIP
5ZVJ        DTDTVIDAIQTDASINHGNAGGPLIDMDAQVIGINTAG--KSLSDSAS-----GLGFAIP
3GDV        NPTGRQNFLQTDASINPGNSGGALVNSLGELMGINTLS--FDKSNDGE--TPEGIGFAIP
7CO3        GLNTYEDFIQTDAAINPGNAGGALVDAAGNLIGINTAI--FSK--SGG---SQGIGFAIP
3PV5        NIEGVENFIQTDAAIGGGNSGGALVNAKGELIGINTAI--LSP--YGG---NVGIGFAIP
4YNN        NIEGVENFIQTDAAINPGNAGGALVNAKGELIGINTAI--ISP--YGG---NVGIGFAIP
4A9G        NLEGLENFIQTDASINRGNAGGALLNLNGELIGINTAI--LAP--GGG---SVGIGFAIP
2ZLE        NAENYENFIQTDAAINRGNSGGALVNLNGELIGINTAI--LAP--DGG---NIGIGFAIP
6JJO        NAENYENFIQTDAAINRGNAGGALVNLNGELIGINTAI--LAP--DGG---NIGIGFAIP
5T69        IPDKRVEFIQTDAAINPGNAGGPLLNARGEVIGINTAI--RAD--------ATGIGFAIP
4IC5        NGKSISEAIQTDADINSGNAGGPLLDSYGHTIGVNTAT--FTRKGSGM---SSGVNFAIP
4IC6        TGVTIGGGIQTDAAINPGNAGGPLLDSKGNLIGINTAI--FTQ--TGT---SAGVGFAIP
3QO6        TGRPIQDVIQTDAAINPGNSGGPLLDSSGTLIGINTAI--YSP--SGA---SSGVGFSIP
                    :* ** *. **:**. .:  .  :*:                   .:.: **

4FLN        TTVVSHFLTD------YERNGKYTGYPCLGVLLQKLENPALR-----ECLKVP---TN-E
5ILB        TTVVSHFLTD------YERNGKYTGFPVLGIEWQKMENPDLR-----KSMGME---SHQK
5JYK        TPVIVHFIQD------YEKHDKYTGFPVLGIEWQKMENPDLR-----KSMGME---SHQK
5FHT        SDRLREFLHRGEKKNSSSGISGSQ-RRYIGVMMLTL-SPSILAELQLREPSFP---DVQH
3NZI        SDKIKKFLTESHDRQ-AKGKAITK-KKYIGIRMMSL-TSSKAKELKDRHRDFP---DVIS
4RI0        SDRITRFLTEFQ----DKQIKDWK-KRFIGIRMRTI-TPSLVDELKASNPDFP---EVSS
6Z05        SNMVKDIAKK------LIEKGKID-RGFLGVTILAL-QGDTK-----KAYKNQ------E
2Z9I        VDQAKRIADE------LISTGKAS-HASLGVQVT---NDKDT-----------------L
5ZVJ        VNEMKLVANS------LIKDGKIV-HPTLGISTRSV-SNAIA-----------------S
3GDV        FQLATKIMDK------LIRDGRVI-RGYIGIGGREI-APLHA-----QGGGID---QL-Q
7CO3        TKLALEVMQS------IIEHGQVI-RGWLGVEVKAL-TPELA-----ESLGLG---ET-A
3PV5        INMVKDVAQQ------IIKFGSIH-RGLMGIFVQHL-TPELA-----QAMGYP---EDFQ
4YNN        INMAKDVAQQ------IIKFGSIH-RGLMGIFVQHL-TPELA-----QSMGYA---EDFQ
4A9G        SNMARTLAQQ------LIDFGEIK-RGLLGIKGTEM-SADIA-----KAFNLD---VQ-R
2ZLE        SNMVKNLTSQ------MVEYGQVK-RGELGIMGTEL-NSELA-----KAMKVD---AQ-R
6JJO        SNMVKNLTSQ------MVEYGQVK-RGELGIMGTEL-NSELA-----KAMKVD----AQR
5T69        IDQAKAIQNT------LAAGGTVP-HPYIGVQMMNI-TVDQA-QQNNRNPNSPFIIPEVD
4IC5        IDTVVRTVPY------LIVYGTAY-RDRLSS------VDKLA------------------
4IC6        SSTVLKIVPQ------LIQFSKVL-RAGINIELA---PDPVA-----NQLNVR------N
3QO6        VDTVGGIVDQ------LVRFGKVT-RPILGIKF----APDQS----VEQLGV-------S
                                        :.

4FLN        GVLVRRVEPTSDASKV-LKE-----------GDVIVSFDDLHVGCEGTVPFRSSERIAFR
5ILB        GVRIRRIEPTAPESQV-LKP-----------SDIILSFDGVNIANDGTVPFRHGERIGFS
5JYK        GVRIRRIEPTAPESQV-LKP-----------SDIILSFDGVNIANDGTVPFRHGERIGFS
5FHT        GVLIHKVILGSPAHRAGLRP-----------GDVILAIGEQMV----------QNAEDVY
3NZI        GAYIIEVIPDTPAEAGGLKE-----------NDVIISINGQSV----------VSANDVS
4RI0        GIYVQEVAPNSPSQRGGIQD-----------GDIIVKVNGRPL----------VDSSELQ
6Z05        GALITDVQKGSSADEAGLKR-----------GDLVTKVNNKVI----------KSPIDLK
2Z9I        GAKIVEVVAGGAAANAGVPK-----------GVVVTKVDDRPI----------NSADALV
5ZVJ        GAQVANVKAGSPAQKGGILE-----------NDVIVKVGNRAV----------ADSDEFV
3GDV        GIVVNEVSPDGPAANAGIQV-----------NDLIISVDNKPA----------ISALETM
7CO3        GIVVAGVYRDGPAARGGLLP-----------GDVILTIDKQEA----------SDGRRSM
3PV5        GALVSQVNPNSPAELAGLKA-----------GDIITQINDTKI----------TQATQVK
4YNN        GALVSQVNQNSPAQLAGLKS-----------GDVIVQINDTKI----------TQATQVK
4A9G        GAFVSEVLPGSGSAKAGVKA-----------GDIITSLNGKPL----------NSFAELR
2ZLE        GAFVSQVLPNSSAAKAGIKA-----------GDVITSLNGKPI----------SSFAALR
6JJO        GAFVSQVLPNSSAAKAGIKA-----------GDVITSLNGKPI----------SSFAALR
5T69        GILVMRVLPGTPAERAGIRR-----------GDVIVAVDGTPI----------SDGARLQ
4IC5        ------------------------------------------------------------
4IC6        GALVLQVPGKSLAEKAGLHPTSRGFAGNIVLGDIIVAVDDKPV----------KNKAELM
3QO6        GVLVLDAPPSGPAGKAGLQSTKRDGYGRLVLGDIITSVNGTKV----------SNGSDLY
```

```
4FLN      YLISQKFAGDIAEIGII-RAGEHKKVQVVLRPRVHLVPYHIDGGQPSYIIVAGLVFTPLS
5ILB      YLISQKYTGDSALVKVL-RNKEILEFNIKLAIHKRLIPAHISGKPPSYFIVAGFVFTTVS
5JYK      YLISQKYTGDSALVKVL-RNKEILEFNIKLAIHKRLIPAHISGKPPSYFIVAGFVFTTVS
5FHT      EAVRT---QSQLAVQIR-RGRETLTLYVTPEVTE------LEHHHHHH------------
3NZI      DVIKR---ESTLNMVVR-RGNEDIMITVIPEEID---PRSLEHHHHHH------------
4RI0      EAVLT---ESPLLLEVR-RGNDDLLFSIAPEVVMG------GHHHHHH------------
6Z05      NYIGTLEIGQKISLSYE-RDGENKQASFILKGEKENPKGVQS--DLIDGLSLRNLDPRLK
2Z9I      AAVRSKAPGATVALTFQDPSGGSRTVQVTLGKAEQ-----LEHHHHHH------------
5ZVJ      VAVRQLAIGQDAPIEVV-REGRHVTLTVKPDPDSTKLAAALEHHHHHH------------
3GDV      AQVAEIRPGSVIPVVVM-RDDKQLTLQVTIQEYPATN-----------------------
7CO3      NQVARTRPGQKISIVVL-RNGQKVNLTAEVGLRPPPAPAPQQKQDGGE------------
3PV5      TTISLLRVGSTVKIIVE-RDNKPLTLSAVVTDIKSHEQKLQSNNPFLYGLALRAFEQESP
4YNN      TTISLLRAGSTAKIKIL-RDNKPLTLDVEVTDIKKHEQKLQSNNPFLYGLALRNFEQESP
4A9G      SRIATTEPGTKVKLGLL-RNGKPLEVEVTLDTSTSSSASAEMITPALEGATLSDGQL---
2ZLE      AQVGTMPVGSKLTLGLL-RDGKQVNVNLELQQSSQNQVDSSSIFNGIEGAEMSN------
6JJO      AQVGTMPVGSKLTLGLL-RDGKQVNVNLELQQSSQNQVDSSSIFNGIEGAEMSN------
5T69      RIVEQAGLNKALKLDLL-RGDRRLSLTVQTAQLRNPTS----------------------
4IC5      --------------------------------------AALEHHHHHH------------
4IC6      KILDEYSVGDKVTLKIK-RGNEDLELKISLEEKSS-----LEHHHHHH------------
3Q06      RILDQCKVGDEVTVEVL-RGDHKEKISVTLEPKPDESAAALEHHHHHH------------

4FLN      EPLIE----EECEDTIGLKLLTKARYSVARFRGEQIVILSQVLANEVNIGYEDMNNQQVL
5ILB      VPYLRSEYGKEYEFDAPVKLLEKHLHAMAQSVDEQLVVVSQVLVSDINIGYEEIVNTQVV
5JYK      VPYLRSEYGKEYEFDAPVKLLEKHLHAMAQSVDEQLVVVSQVLVSDINIGYEEIVNTQVV
5FHT      ------------------------------------------------------------
3NZI      ------------------------------------------------------------
4RI0      ------------------------------------------------------------
6Z05      DRLQI-------------------------PKDVNGVLVDSVKEKSKGKNSGFQEGDIII
2Z9I      ------------------------------------------------------------
5ZVJ      ------------------------------------------------------------
3GDV      ------------------------------------------------------------
7CO3      ------------------------------------------------------------
3PV5      P-----------------------------HGNVIGVQVVGASENSAGWRAGIRPGDIII
4YNN      P-----------------------------HGNVVGVQVVGASETSAGWRAGLRPGDIII
4A9G      ------------------------------KDGGKGIKIDEVVKGSPAAQAGLQKDDVII
2ZLE      ------------------------------KGKDQGVVVNNVKTGTPAAQIGLKKGDVII
6JJO      ------------------------------KGKDQGVVVNNVKTGTPAAQIGLKKGDVII
5T69      ------------------------------------------------------------
4IC5      ------------------------------------------------------------
4IC6      ------------------------------------------------------------
3Q06      ------------------------------------------------------------

4FLN      KFNGIPIRNIHHLAHLIDMCKDKYLVFEFEDNYVAVLEREASNSASLCILKDYGIPSERS
5ILB      AFNGKPVKNLKGLAGMVENCEDEYMKFNLDYDQIVVLDTKTAKEATLDILTTHCIPSAMS
5JYK      AFNGKPVKNLKGLAGMVENCEDEYMKFNLDYDQIVVLDTKTAKEATLDILTTHCIPSAMS
5FHT      ------------------------------------------------------------
3NZI      ------------------------------------------------------------
4RI0      ------------------------------------------------------------
6Z05      GVGQSEIKNLKDLEQALKQVNKKEFTKVWVYRNGFATLLVLK------------------
2Z9I      ------------------------------------------------------------
5ZVJ      ------------------------------------------------------------
3GDV      ------------------------------------------------------------
7CO3      ------------------------------------------------------------
3PV5      SANKKPVTDVKSLQTIAQEKKKELLVQVLRGPGSMYLLVI--------------------
4YNN      SANKTPVKDIKSLQAVAHEKGKQLLVQVLRGAGALYLLII--------------------
4A9G      GVNRDRVNSIAEMRKVLAAKPAIIALQIVRGNESIYLLMRLEHHHHHH------------
2ZLE      GANQQAVKNIAELRKVLDSKPSVLALNIQRGDSTIYLLMQ--------------------
6JJO      GANQQAVKNIAELRKVLDSKPSVLALNIQRGDSTIYLLMQ--------------------
5T69      ------------------------------------------------------------
4IC5      ------------------------------------------------------------
4IC6      ------------------------------------------------------------
3Q06      ------------------------------------------------------------
```

```
4FLN            ADLLEPYVDPIDDTQALDQGIGDSPVSNLEIGFDGLVWA
5ILB            DDLKTEERN------------------------------
5JYK            DDLKTEERN------------------------------
5FHT            ---------------------------------------
3NZI            ---------------------------------------
4RI0            ---------------------------------------
6Z05            ---------------------------------------
2Z9I            ---------------------------------------
5ZVJ            ---------------------------------------
3GDV            ---------------------------------------
7CO3            ---------------------------------------
3PV5            ---------------------------------------
4YNN            ---------------------------------------
4A9G            ---------------------------------------
2ZLE            ---------------------------------------
6JJO            ---------------------------------------
5T69            ---------------------------------------
4IC5            ---------------------------------------
4IC6            ---------------------------------------
3QO6            ---------------------------------------
```

**<u>SI Figure 5:</u> Secondary Structures of the HtrA modules**

Cartoon representation of the secondary structures of β-barrel modules in the HtrA proteases. A) Arrangement of the strands in the protease domain modules. Strands are colored from N- to C-terminus orange, yellow, green, cyan, blue, and magenta respectively. B) Arrangement of the strands in the PDZ-like domain modules. The PDZ-like domain modules have four strands that correspond to the same spatial positions in the protease domain modules and those strands are colored their equivalent color. There are no equivalent strands in the green or magenta positions in the PDZ-like modules. Additionally, the first strand in the sequence which has been rotated out of the barrel is colored in grey. A sixth strand is present in some of the proteins, but it too has also been rotated out of the barrel and is colored in grey in the figure as well. Note that the alternating, antiparallel arrangement of the strands is maintained in the PDZ-like domains, even if it is formally reversed (as the definition of positive and negative is an arbitrary choice).

A B

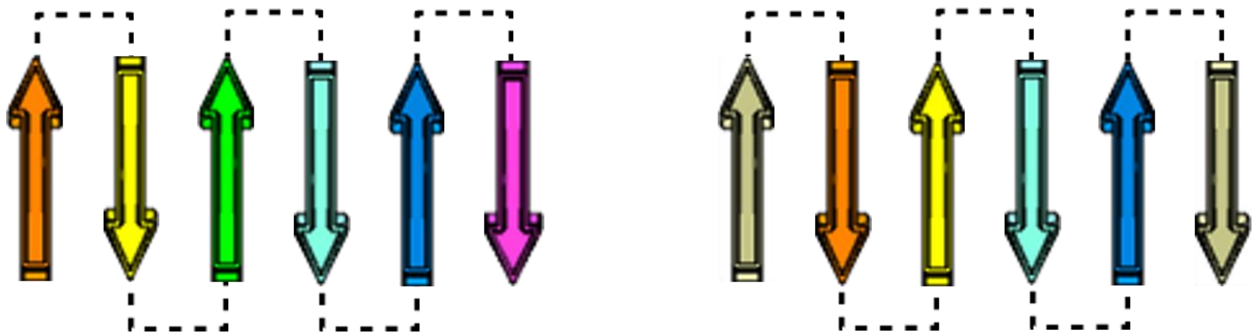

**<u>SI Figure 6</u>: Structures of the HtrA modules**

Cartoon representation of the modules of the sequence unique HtrA protease structures showing the alpha helices (red) and beta strands (yellow). Coil regions are hidden to simplify the images. Figures made in PyMol. A) All the N-terminal protease modules B) All the C-terminal protease modules C) All the PDZ modules D) All modules (both protease and PDZ) E) All three modules from (PDB ID 5t69) with the individual strands colored based on their structural position with sequence shown and highlighted (colors from N to C in the protease domain: orange, yellow, green, blue, and magenta).

A)

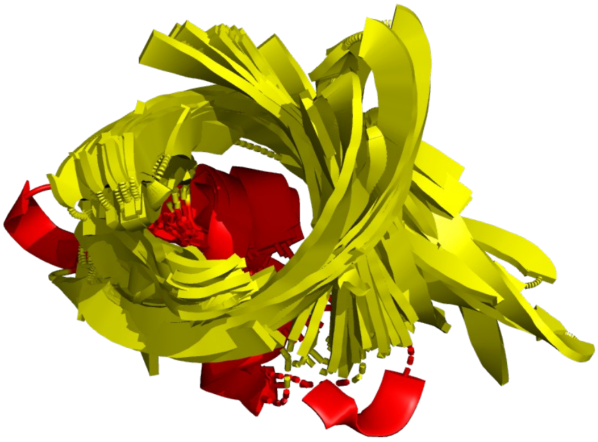

B)

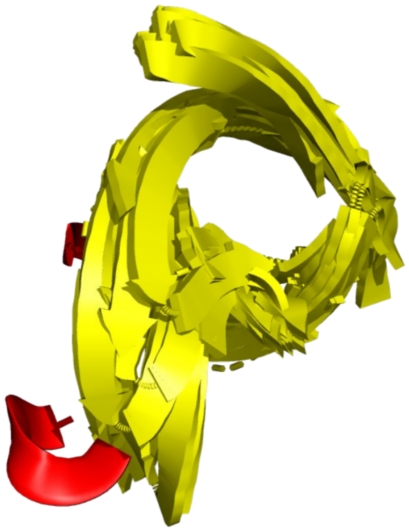

C)

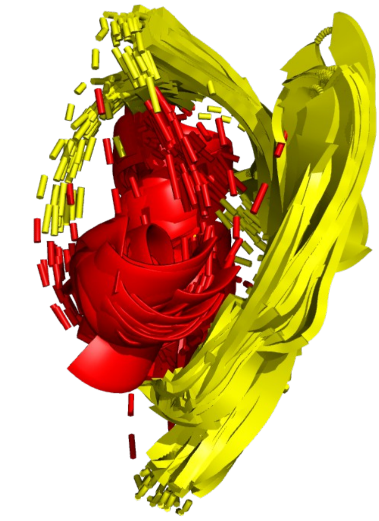

D)

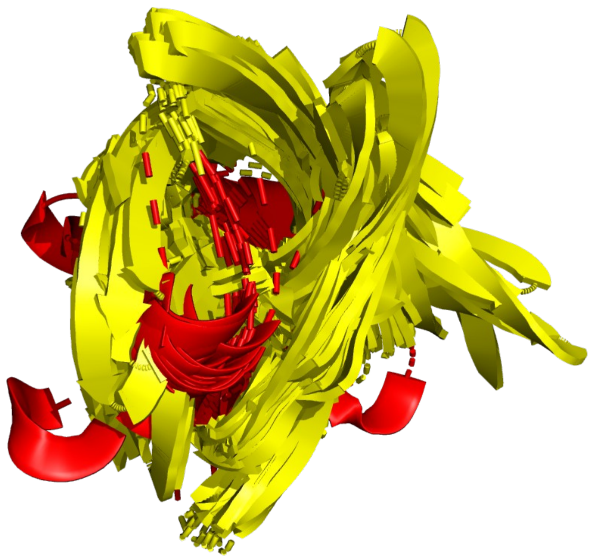

E)

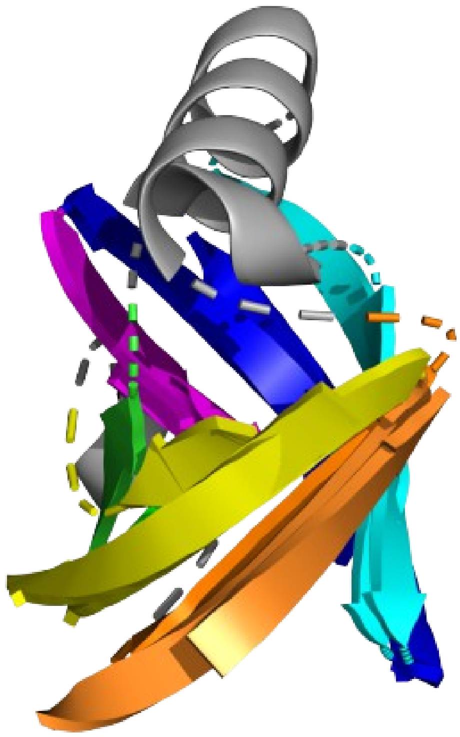

**<u>SI Table 1:</u> Domain Sequence Similarity Comparision**

Comparision of the average sequence identity between the different domains in the HtrA protease PDB structure set. Identity was calculated by pairwise comparison using Needleman-Wunsch alignments. Domains were defined using the PDB structures.

| % ID (mean) | Protease | PDZ1 | PDZ2 |
|---|---|---|---|
| Protease | 20.1 | | |
| PDZ1 | 18.2 | 21.2 | |
| PDZ2 | 16.4 | 19.9 | 26.1 |

| % ID (median) | Protease | PDZ1 | PDZ2 |
|---|---|---|---|
| Protease | 15.4 | | |
| PDZ1 | 15.3 | 15.5 | |
| PDZ2 | 14.2 | 16.3 | 17.1 |

**SI Table 2: Unmodified RMSD comparison table**

RMSD calculated (CE-align in PyMol) comparing modules from a given protein structure to its own other modules

| Protein | Prot1-Prot2 | Prot1-PDZ1 | Prot2-PDZ1 | Prot1-PDZ2 | Prot2-PDZ2 | PDZ1-PDZ2 |
|---|---|---|---|---|---|---|
| 2zle | 4.671412 | 5.393964 | 9.871169 | 9.197718 | 8.582769 | 2.893496 |
| 2z9i | 4.645298 | 6.99783 | 9.945109 | | | |
| 3gdv | 4.560794 | 10.308235 | 8.325793 | | | |
| 3nzi | 4.203474 | | | | | |
| 3pv5 | 3.27669 | 9.093188 | 9.69562 | | | |
| 3qo6 | 3.941116 | 9.468741 | 9.208978 | | | |
| 4a9g | 4.129794 | 9.667325 | 9.413048 | 7.615468 | 9.794967 | 4.521349 |
| 4fln | 3.559123 | 9.572902 | 9.204368 | 10.260514 | 10.506852 | 4.814594 |
| 4ic5 | 5.256993 | | | | | |
| 4ic6 | 4.429048 | 8.826231 | 9.802585 | | | |
| 4ri0 | 5.019885 | 7.696705 | 6.892202 | | | |
| 4ynn | 3.398238 | 9.181668 | 10.269787 | 10.343585 | 9.604559 | 2.40976 |
| 5fht | 4.189576 | 9.519808 | 9.579156 | | | |
| 5ilb | 3.420169 | 8.852867 | 9.15623 | 10.051552 | 7.966859 | 4.95404 |
| 5jyk | 3.094949 | 9.052668 | 9.531856 | 8.316712 | 11.163155 | 5.449278 |
| 5t69 | 2.748297 | 10.537871 | 5.853515 | | | |
| 5zvj | 4.54295 | 5.341272 | 9.874509 | | | |
| 6jjo | 2.862183 | 9.128376 | 9.484773 | 9.207652 | 7.006915 | 2.177693 |
| 6z05 | 6.0494 | 9.362718 | 10.224059 | 8.803575 | 9.93282 | 4.125754 |
| 7co3 | 3.198494 | 9.784797 | 9.943692 | | | |
| **Mean =** | 4.05989415 | 8.76595367 | 9.2375805 | 9.224597 | 9.319862 | 3.9182455 |
| **Median =** | 4.159685 | 9.155022 | 9.555506 | 9.202685 | 9.699763 | 4.3235515 |

**SI Table 3:** **β-barrel comparison summary**

Summary table for comparisons of the β-barrel modules in the HtrA proteases. "all" indicates that pairwise comparisons of all modules were included to calculate the comparison value. "self" indicates only pairwise comparisons between protease or between PDZ modules were included. "other" indicates pairwise comparisons of protease domains to PDZ domains were used to calculate the comparison value. "proteases" indicates only comparisons between protease modules were used. "PDZ" indicates that only comparison between PDZ modules were used to calculate the comparison values. "PDZ1" indicates only comparisons to the first PDZ domain were used.

**CEALIGN to self (RMSD, Å)**

| | Proteases | Proteases to PDZ1 |
|---|---|---|
| **Mean =** | 4.060 | 9.002 |
| **Median =** | 4.160 | 9.441 |

**CEALIGN loops removed (RMSD, Å)**

| | All | Self | Other |
|---|---|---|---|
| **Mean =** | 6.444 | 5.392 | 7.056 |
| **Median =** | 6.789 | 5.002 | 7.460 |

**CEALIGN to 5m3n (RMSD, Å)**

| | N-protease | C-protease | PDZ1 |
|---|---|---|---|
| **Mean =** | 2.890 | 2.936 | 2.124 |
| **Median =** | 2.500 | 2.819 | 1.677 |

**TM Align (RMSD, Å)**

| | Proteases | PDZ | Other |
|---|---|---|---|
| **Mean =** | 2.804 | 2.273 | 3.724 |
| **Median =** | 2.780 | 2.215 | 3.695 |

**TM Align (TM score)**

| | Proteases | PDZ | Other |
|---|---|---|---|
| **Mean =** | 0.552 | 0.552 | 0.298 |
| **Median =** | 0.549 | 0.549 | 0.299 |
